# Epiregulon: Inference of single-cell transcription factor activity to predict drug response and drivers of cell state

**DOI:** 10.1101/2023.11.27.568955

**Authors:** Tomasz Włodarczyk, Aaron Lun, Diana Wu, Minyi Shi, Xiaofen Ye, Shreya Menon, Shushan Toneyan, Kerstin Seidel, Liang Wang, Jenille Tan, Shang-Yang Chen, Timothy Keyes, Aleksander Chlebowski, Adrian Waddell, Wei Zhou, Yangmeng Wang, Qiuyue Yuan, Yu Guo, Liang-Fu Chen, Bence Daniel, Antonina Hafner, Meng He, Alejandro Chibly, Yuxin Liang, Zhana Duren, Ciara Metcalfe, Marc Hafner, Christian W. Siebel, M. Ryan Corces, Robert Yauch, Shiqi Xie, Xiaosai Yao

## Abstract

Transcription factors (TFs) and transcriptional coregulators represent an emerging class of therapeutic targets in oncology. Gene regulatory networks (GRNs) can be used to evaluate pharmacological agents targeting these factors and to identify drivers of disease and drug resistance. However, GRN methods that rely solely on gene expression often fail to account for post-transcriptional modulation of TF function. We present Epiregulon, a method that constructs GRNs from single-cell ATAC-seq and RNA-seq data for accurate prediction of TF activity. This is achieved by considering the co-occurrence of TF expression and chromatin accessibility at TF binding sites in each cell. We leverage ChIP-seq data to extend inference to transcriptional coregulators lacking defined motifs or TF harboring neomorphic mutations. Epiregulon accurately predicted the effects of AR inhibition across various drug modalities including an AR antagonist and an AR degrader, delineated the mechanisms of a SMARCA4 degrader by identifying context-dependent interaction partners and prioritized known and novel drivers of lineage reprogramming and tumorigenesis. By mapping gene regulation across various cellular contexts, Epiregulon can accelerate the discovery of therapeutics targeting transcriptional regulators.

## Introduction

Transcription factors and transcriptional coregulators shape cell fates and lineage commitment, and their dysregulation drives congenital diseases and tumor growth. TFs bind to specific DNA sequences whereas coregulators interact with TFs in a context-specific manner and lack defined motifs. These transcriptional modulators represent an important class of therapeutic targets in oncology and beyond. Current therapeutics targeting transcriptional regulators inhibit their activity by either blocking the ligand binding domains^1^, degrading the protein^2,3^ or disrupting protein-protein interactions^4^. Delineating the functions of transcriptional regulators can accelerate our understanding of disease biology and drug discovery.

Gene regulatory networks (GRNs) model the underlying circuitry of gene regulation and have been applied to understand lineage commitment, plasticity and drug response^5^. Early methods to construct gene regulatory networks relied exclusively on gene expression data and sought to identify association between expression of TFs and their target genes^6,7^. Now, single-cell multiomics technologies provide chromatin accessibility information in addition to gene expression and recent GRN inference methods leverage both modalities to improve performance. Most of these methods, including CellOracle^8^, FigR^9^, Pando^8^ and GRaNIE^10^, rely on linear relationships between the mRNA levels of transcription factors and their target genes to model gene regulation. In addition, SCENIC+ utilizes random forest regression to model the regulatory relationships between TFs, regulatory elements and target genes^11^.

Several challenges still impede the broader utilization of these GRN inference methods in basic biology and drug discovery. First, none of these methods were specifically designed to predict changes in which the TF activity is decoupled from its gene expression. These include drug perturbations that disrupt protein complex formation or localization and post-translational modifications that can impact TF activity and genetic alterations (e.g. neomorphic mutations and CRISPR genome editing) that can silence TF function or add new functions without changing gene expression. Second, the use of motif sequences in GRN construction precludes activity inference of transcriptional coregulators, an important class of regulators without defined motifs. This is problematic for methods that use motifs to filter target genes based on putative TF binding sites.

Here we present Epiregulon, a method that constructs GRNs from single-cell multiomics and TF occupancy data to infer activity of transcriptional regulators. Epiregulon uses the co-occurrence of TF mRNA expression and chromatin accessibility at TF binding sites to accurately determine the relevance of potential target genes in any given biological context. Unlike most other GRN tools, Epiregulon also leverages ChIP-seq data to infer the activity of TFs and transcriptional coregulators lacking defined motifs. Functional interpretation of the built GRN is performed by computing the Jaccard similarity of target genes and known pathway annotations. We applied Epiregulon to several real and simulated datasets where it successfully recovered the ground truth, detected novel regulators, and matched or out-performed existing GRN methods. Our results indicate that Epiregulon is well suited to infer single-cell TF activity in the context of drug perturbation and lineage reprogramming. Epiregulon is implemented as a suite of open-source R packages that are available from the Bioconductor project.

## Results

### Epiregulon constructs GRNs to infer TF activity at the single cell level

Epiregulon is designed to infer the activity of a transcription factor (TF) or a transcriptional coregulator (collectively referred to as TFs for brevity) under a variety of biological scenarios: 1) regulator activity driven by overexpression, 2) regulator activity decoupled from mRNA expression, 3) a context-dependent co-regulator interacting with different TFs and 4) gain of function due to neomorphic mutations or hijacking by other factors (**Figure 1a**).

**Figure 1.**
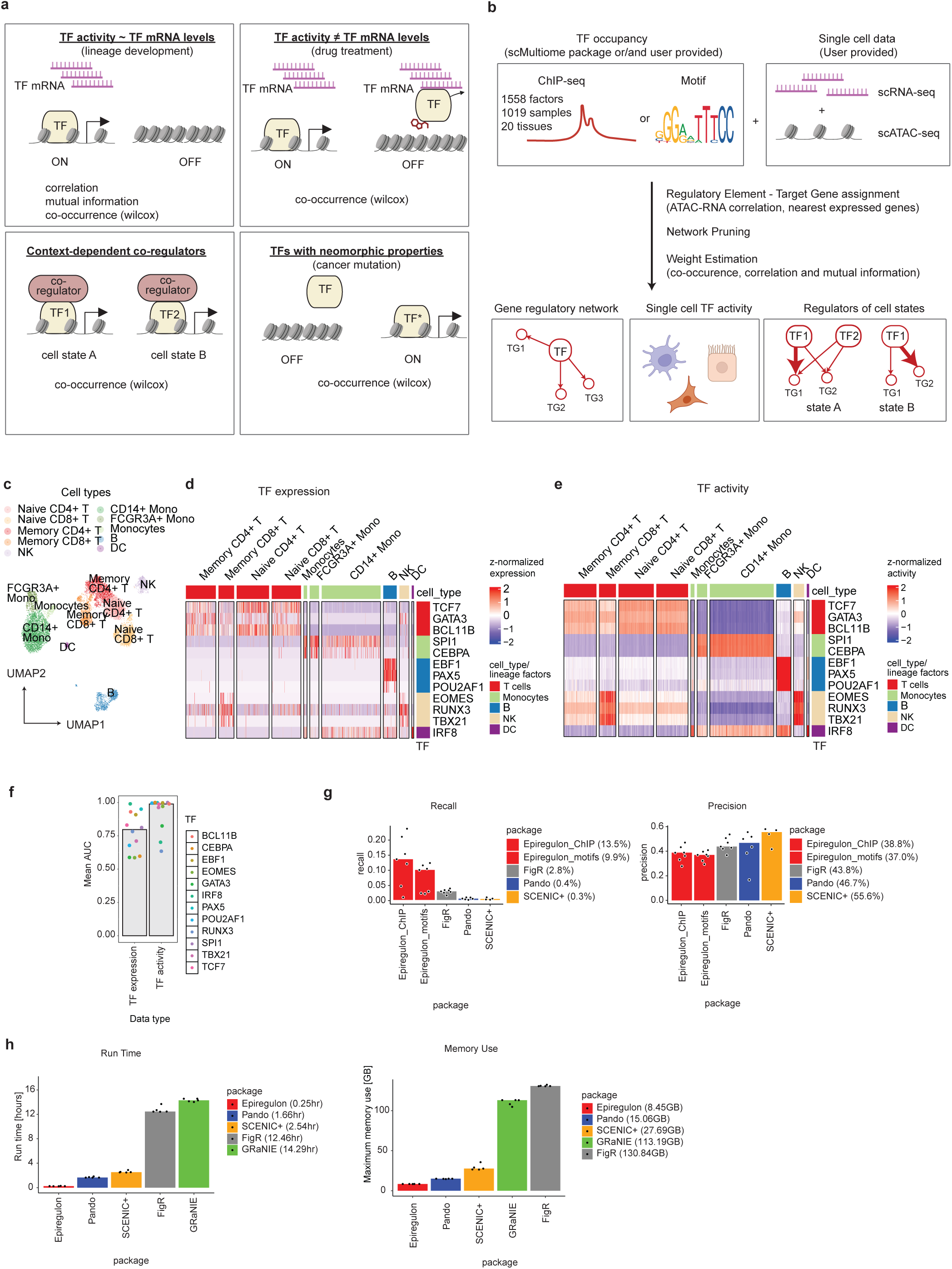
Epiregulon constructs GRNs to infer regulator activity at the single cell level. a) Epiregulon can infer regulator activity in a multitude of scenarios. The first scenario is when the activity of a transcription factor is regulated by its gene expression such as in lineage development. The second scenario is when TF activity is discordant with its mRNA expression as in the case of TF inhibition by a drug molecule. Thirdly, Epiregulon can estimate the activity of a motif-lacking co-regulator that may associate with different TFs depending on cell context. Lastly, Epiregulon can estimate regulators harboring neomorphic mutations using pan-cell-type binding sites or user-provided ChIP-seq data. Correlation and mutual information weight estimation methods are appropriate for the first scenario whereas co-occurrence is applicable to all cases. b) If users provide scRNA-seq and scATAC-seq, Epiregulon can construct GRNs either from ChIP-seq data or motif annotations. Pan-cell-type, tissue-specific and sample-specific ChIP-seq data compiled from ChIP-Atlas and ENCODE are available through the scMultiome package. Epiregulon outputs regulator activity at the single cell level, a pruned and weighted gene regulatory network and differential activity analysis to identify potential drivers of cell states. c) For benchmarking, we downloaded the paired scATAC-seq and scRNA-seq PBMC data from 10x genomics. We identified cell types using SingleR and known marker genes. Shown is the UMAP representation of the various cell types present in the data. d) Gene expression of known lineage markers e) Activities of the same lineage markers were calculated using Epiregulon (correlation weight estimation method). f) Area under the receiver operating characteristics curve (AUROC) is calculated based on whether TF expression or TF activity can distinguish cells of the matching lineage vs the remaining cells. g) Gene expression changes after depletion of 7 individual factors (ELK1, GATA3, JUN, NFATC3, NFKB1, STAT3 and MAF) were obtained from the knockTF database and genes with absolute logFC > 0.5 and corrected p-value < 0.05 were considered ground truth target genes. GRNs obtained from the shown packages were evaluated for precision and recall of target genes. h) Run time and memory use of the GRN construction from the PBMC data were evaluated with 64GB and 20 cores on high performance computing (HPC) cluster. In the case of GRaNIE the memory allocation needs to be increased to 128 GB and for FigR the memory allocation was increased to 256 GB. Each package was run 5 times.

Epiregulon infers TF activity from a single-cell multiomics dataset containing paired RNA-seq and ATAC-seq counts (**Figure 1b**). The ATAC-seq data is first used to identify regulatory elements (REs) from regions of open chromatin. The REs are filtered to those that overlap the binding sites of the TF, typically determined from external ChIP-seq data. Epiregulon provides a pre-compiled list of ChIP-seq binding sites from ENCODE and ChIP-Atlas spanning 1558 factors, 1019 cell types/lines and 20 tissues (refer to **Supplementary Table S1** and methods for statistics and quality control). This list of sites can be further stratified by cell line or tissue, depending on the user’s biological context. We also provide options to identify sites from motif annotations (see methods, **Figure 1b**) or to use user-supplied binding sites.

Once a list of relevant REs has been obtained, each RE is tentatively assigned to all genes within a distance threshold. A gene is considered a target gene (TG) if the correlation between ATAC-seq and RNA-seq counts across cells in the paired single-cell data is strong. Each RE-TG edge is then assigned a weight using the “co-occurrence method”, defined as the Wilcoxon test statistic from the comparison of the TG expression between “active” cells (that both express the TF and have open chromatin at the RE) to all other cells. The co-occurrence method is the default weighting scheme as it is less reliant on the degree of TF expression and thus can handle situations where TF activity is decoupled from expression. However, other weights can be used for TFs where activity is driven by expression (e.g. correlation or mutual information between TF and TG expression). Simulations indicate that Epiregulon’s results are robust to the choice of weighting scheme, even with sparse data and false connections (**Supplementary Note Section 1**).

After repeating the above steps for all TFs of interest, we obtain a weighted tripartite graph spanning the TFs, the REs overlapping their binding sites and the neighboring TGs. This weighted graph is the final GRN that is returned by Epiregulon. For each cell, the predicted activity of a TF is defined as the RE-TG-edge-weighted sum of the expression values of its TGs in the GRN, divided by the number of TGs. Epiregulon can also test for differential activity between conditions via total activity or edge subtraction of the GRN (**Supplementary Note Section 2)** to identify potential context-dependent interaction partners of each TF. A more detailed description of the entire Epiregulon algorithm is provided in the Methods and **Figure S1**.

### Epiregulon yields a high recall of target genes in PBMC data

We first evaluated the performance of Epiregulon using a human peripheral blood mononuclear cell (PBMCs) dataset obtained from 10x Genomics (**Figure 1c**). The estimated activities of known lineage factors aligned to their respective lineages with greater specificity compared to TF expression alone (**Figure 1d-f**). These lineage factors include TCF7, GATA3 and BCL11B (T cells), SPI1 and CEBPA (myeloid cells), EBF1, PAX5 and POU2AF1 (B cells), EOMES, RUNX3 and TBX21 (NK cells) and IRF8 (dendritic cells, DC). Epiregulon revealed that, in addition to being a lineage factor of NK cells, TBX21 exhibited heightened activity in CD8+ memory T cells, consistent with the depletion of this cell type in *Tbx21*^−/−^ mice^12^. IRF8 activity was also elevated in DCs and moderate in monocytes, consistent with its known functions in myeloid development^13^.

Next, we benchmarked Epiregulon and other GRN methods (**Supplementary Table S2)** based on their ability to predict target genes for a comparison of GRN methods. From the knockTF database, we identified 7 factors that were depleted in human blood cells - ELK1, GATA3, JUN, NFATC3, NFKB1, STAT3 and MAF. Genes with altered expression upon depletion of each TF were considered true target genes of that TF. Epiregulon detected more of these altered genes in the PBMC dataset than other GRN methods, at the cost of a modest loss in precision (**Figure 1g, Figure S2**). SCENIC+ was the most precise method but failed to return a GRN for 3 of the 7 lineage factors (**Figure 1g**). These results indicate that each method achieves a different compromise between recall and precision and Epiregulon is most suited for high recovery of target genes. Epiregulon also used the least computational time and memory (**Figure 1h**), which is advantageous for iterative analyses.

### Epiregulon predicts the responses of AR-modulating drugs

A more challenging task is to predict changes when TF activity is decoupled from its gene expression. We generated a single-cell multiomics dataset to evaluate changes in AR activity following drug treatment. Six prostate cancer cell lines (4 AR-dependent and 2 AR-independent) were treated with 3 AR-modulating agents (**Figure 2a**, **2b**). The first agent is the clinically approved AR antagonist, enzalutamide, which interferes with AR protein function by blocking its ligand binding domain^1^. The second agent is ARV-110, a degrader of AR protein which acts by bringing an E3 ubiquitin ligase in close proximity to an AR protein^3^ (**Figure 2c**). We also synthesized a third agent, SMARCA2_4.1, a degrader of SMARCA2 and SMARCA4, two mutually exclusive paralog proteins that encode the ATPase subunit of the SWI/SNF chromatin remodeler, which is crucial for recruiting AR to the chromatin^14^ (**Figure 2d**). These pharmacological agents do not directly or consistently suppress AR mRNA levels.

**Figure 2:**
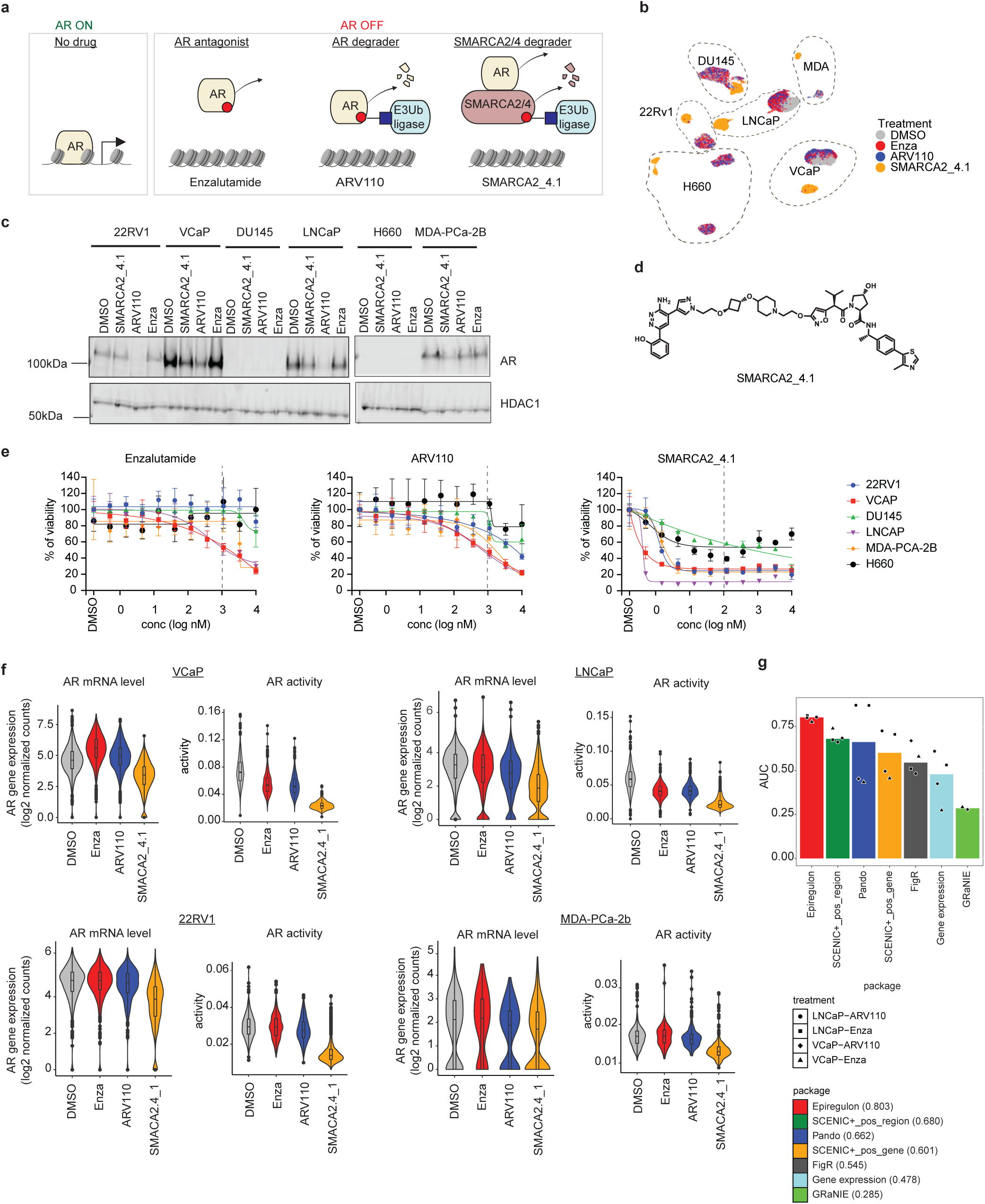
Epiregulon predicts the responses of AR-modulating drugs. a) We investigate AR activity after perturbation with 3 different AR-modulating agents. Enzalutamide is an AR antagonist that competes with androgen for the ligand binding domain of AR. ARV110 is a bifunctional molecule that bridges AR with an E3 ubiquitin ligase for degradation. SMARCA2_4.1 is a degrader of SMARCA2/4, the ATPase subunit of the SWI/SNF remodeling complex that is required for the binding of AR to the chromatin. b) Six prostate cancer cell lines were treated with the 3 AR-modulating agents and profiled for changes in their gene expression and chromatin accessibility by paired scATAC-seq and scRNAseq. VCaP, LNCaP, 22Rv1 and MDA-PCa-2b are AR-dependent whereas DU145 and H660 are AR-independent. Shown is the UMAP representation (5028 VCaP cells, 5958 LNCaP cells, 3568 22Rv1 cells, 945 MDA-PCa-2b cells, 3639 NCI-H660 cells and 3980 DU145 cells). Cells were merged from 2 technical replicates. Cells were treated for 24 hours at 1 uM of enzalutamide or ARV110 or 0.1 uM of SMARCA2_4.1. c) Immunoblotting of AR and HDAC as a loading control after 24 hours of treatment. d) Chemical structure of SMARCA2_4.1, a degrader of SMARCA2 and SMARCA4 e) Prostate cancer cell lines were treated at varying doses for 5 days and cell viability was measured by CellTiter-Glo. Dotted line indicates the concentrations used in the scATAC-seq/scRNA-seq experiment. f) Shown are the AR gene expression and the AR activity computed by Epiregulon (co-occurrence weight estimation method) g) Each cell was identified by the HTO tag corresponding to the well receiving DMSO or one of the two known AR inhibitors (enzalutamide or ARV110) and this information served as the ground truth. A cell was classified into either treated with DMSO control or an AR inhibitor based on AR activity. AUROC was computed for all the packages being benchmarked as well as for AR gene expression. Bar plots show the median AUROCs in the two sensitive cell lines LNCaP and VCaP treated with enzalutamide or ARV110.

We measured the response of all 6 cell lines to these pharmaceutical targets using cellTiterGlo at 1 and 5 days of treatment. At 1 day post treatment, there was minimal cell death **(Figure S3a**), allowing us to profile their gene expression and chromatin changes. After 5 days of treatment, LNCaP and VCaP exhibited substantial cell death after enzalutamide treatment but MDA-PCa-2b and 22RV1 remained resistant (**Figure 2e)**. Similarly, LNCaP and VCaP showed the greatest sensitivity towards AR degrader ARV110 while MDA-Pca-2b and 22RV1 were mildly responsive. Neither of the AR-independent cell lines responded to enzalutamide or ARV110. All 6 cell lines responded to SMARCA2_4.1 treatment (**Figure 2e**).

We used Epiregulon to predict changes in AR activity as a result of drug treatment. We leveraged publicly available AR ChIP-seq data for LNCaP, VCaP and 22Rv1 and generated our own ChIP-seq for MDA-PCa-2b. Consistent with the observed drug efficacy, Epiregulon predicted decreased AR activity following enzalutamide and ARV110 treatment in the known sensitive cell lines LNCaP and VCaP, and minimal changes in AR activity in the resistant cell lines 22Rv1 and MDA-Pca-2b (**Figure 2f**). In particular, Epiregulon predicted AR activity correctly in VCaP cells despite discordant trends in AR expression (**Figure 2f**), highlighting the utility of the GRN approach.

We benchmarked Epiregulon against other GRN inference methods based on the ability of each method’s AR activity estimates to discriminate between DMSO- and drug-treated cells. Epiregulon was the most accurate method at distinguishing control and treated cells in the sensitive cell lines (**Figure 2g, Figure S3b**). Pando was comparably accurate in LNCaP but performed poorly in VCaP (**Figure S3b**); this is in part because enzalutamide treatment increased AR expression in VCaP cells despite AR inhibition (**Figure 2f).** Lack of response to enzalutamide and slight response to ARV110 were observed for the resistant cell lines MDA-PCa-2b and 22Rv1 and (**Figure S3c**). These results indicate that Epiregulon should be the method of choice to predict drug efficacy from regulator activity.

### Epiregulon estimates activity of AR harboring neomorphic mutations

An obvious alternative method to quantify AR activity is to compute gene set scores for existing AR signatures^15–19^. This assumes that the GRN of our biological system of interest is similar to that of the system from which the signatures were derived. For example, signatures work well in VCaP (**Figure 3a-b**), likely because amplification of wildtype AR is a frequent event in AR-dependent tumors^15,19^. In contrast, the MDA-PCa-2b cell line harbors two mutations (L702H, T787A) in AR that changes its ligand specificity (**Figure S4a**) and the combination of these 2 mutations is well represented in patient tumors. We hypothesized that these mutations would also alter AR’s regulatory behavior, and indeed, many canonical AR targets such as KLK3 and TMPRSS2 were not suppressed by any of the AR modulating agents **(Figure 3c-d)**. The change in AR function compromises the use of existing signatures, all of which failed to predict a decrease in AR activity upon treatment (**Figure S4b, Supplementary Table S3**).

**Figure 3:**
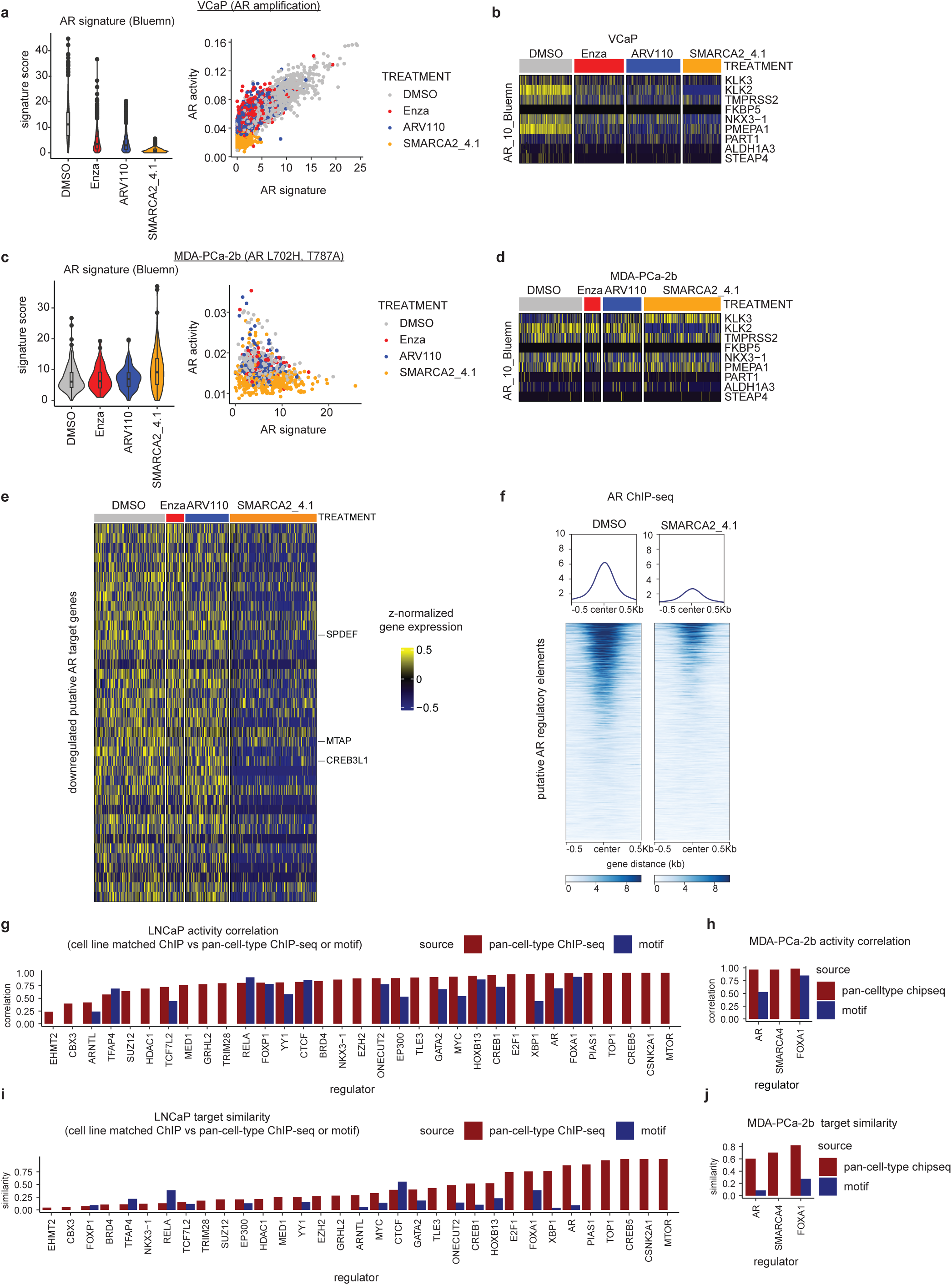
Epiregulon infers activity of AR harboring neomorphic mutations. a) Shown are the AR activity estimated from the signature score in Bluemn et al^15^ and its correlation with AR activity estimated by Epiregulon in VCaP cells which harbor an amplification of the wildtype AR. b) Normalized expression of genes in the Bluemn AR signature for VCaP c) Same as a) but for MDA-PCa-2b which harbors two mutations in the AR gene and as a result has enhanced specificity for hydrocortisone over 5a-DHT. d) Same as b) but for MDA-PCa-2b e) Normalized expression of AR targets of MDA-PCa-2b as inferred by Epiregulon f) AR occupancy as measured by ChIP-seq at ATAC-peaks containing the regulatory elements mapped to AR target genes in MDA-PCa-2b cells treated with DMSO or SMARCA2/4 degrader, SMARCA2_4.1. Center represents the center of the regulatory elements. g) The ground truth regulator activity was computed using all the publicly available ChIP-seq obtained in LNCaP cells. This activity was then correlated (Pearson’s) with activity computed either from pan-cell-type ChIP-seq (red) or motif annotations (blue) for each of the regulators. h) Same as g) but using ChIP-seq generated in MDA-PCa-2b i) Shown is the Jaccard similarity between the target genes derived from LNCaP ChIP-seq vs target genes derived from pan-cell-type ChIP-seq (red) or motif annotations (blue) j) Same as h) but using ChIP-seq generated in MDA-PCa-2b

We previously observed that Epiregulon predicted a decrease in AR activity after SMARCA2_4.1 treatment in MDA-PCa-2b cells (**Figure 2f**). Further investigation of this result revealed that Epiregulon predicted a different set of AR targets for MDA-PCa-2b compared to VCaP, many of which were downregulated by SMARCA2_4.1 treatment (**Figure 3e, Figure S4c-d**). ChIP-seq confirmed reduced AR occupancy at the regulatory elements of AR target genes upon treatment (**Figure 3f**), suggesting that Epiregulon’s inferred AR targets are more accurate than the canonical signatures. This explains the improved performance of Epiregulon over signature scores at predicting drug response in the presence of neomorphic mutations.

### Pan-cell-type ChIP-seq outperforms motifs for accurate estimation of TF activity

Ideally, ChIP-seq data is available for the system of interest as this provides the most accurate information about the TF’s binding sites. In the absence of such data, Epiregulon’s precompiled list of pan-cell-type ChIP-seq binding sites allows users to use information from other cell types, cell lines or tissues for exploratory analysis. Activities predicted by Epiregulon using the pan-cell-type binding sites were highly correlated with the activities obtained using the ideal cell -line-matched ChIP-seq data (**Figure 3g-h**, **Figure S4e-f**). In contrast, activities predicted by motif annotations did not correlate well with cell-line-matched ChIP-seq (**Figure 3g-h**, **Figure S4e-f**).), and many factors did not even have motif annotations. These results suggest that even unmatched ChIP-seq data should be generally preferred to motif annotations for estimating TF activity.

To further investigate the performance of Epiregulon with the pan-cell-type list, we examined the degree of overlap between the target genes identified by Epiregulon with the pan-cell-type sites and those identified with cell-line-matched ChIP-seq. We observed strong overlap for several factors in multiple cell lines (**Figure 3i**-**j****, Figure S4g**), indicating that the pan-cell-type list can often be good enough for target gene identification when cell-line-matched ChIP-seq data is not available. However, other factors exhibited a weaker overlap (**Figure 3i**-**j****, Figure S4g**), suggesting that cell-line-matched ChIP-seq can still be beneficial for pinpointing specific target genes.

### Epiregulon uncovered context-dependent effects of SMARCA4 degradation

SMARCA4 is a transcriptional coregulator that is responsible for proliferation of prostate cancer cell lines^20^. SMARCA2_4.1 effectively depleted SMARCA4 protein expression in all 6 cell lines at 24 hours (**Figure 4a**), so we would expect to see a decrease in SMARCA2_4.1 activity from each GRN method. However, SMARCA4 does not have a well-defined motif as it can interact with different TFs depending on the cellular context. This precludes the use of existing methods that rely on motif annotations for GRN construction. In contrast, Epiregulon can use public SMARCA4 ChIP-seq data to determine the most likely target genes without relying on motifs, upon which it correctly predicts decreased SMARCA4 activity in all cell lines (**Figure 4b**).

**Figure 4:**
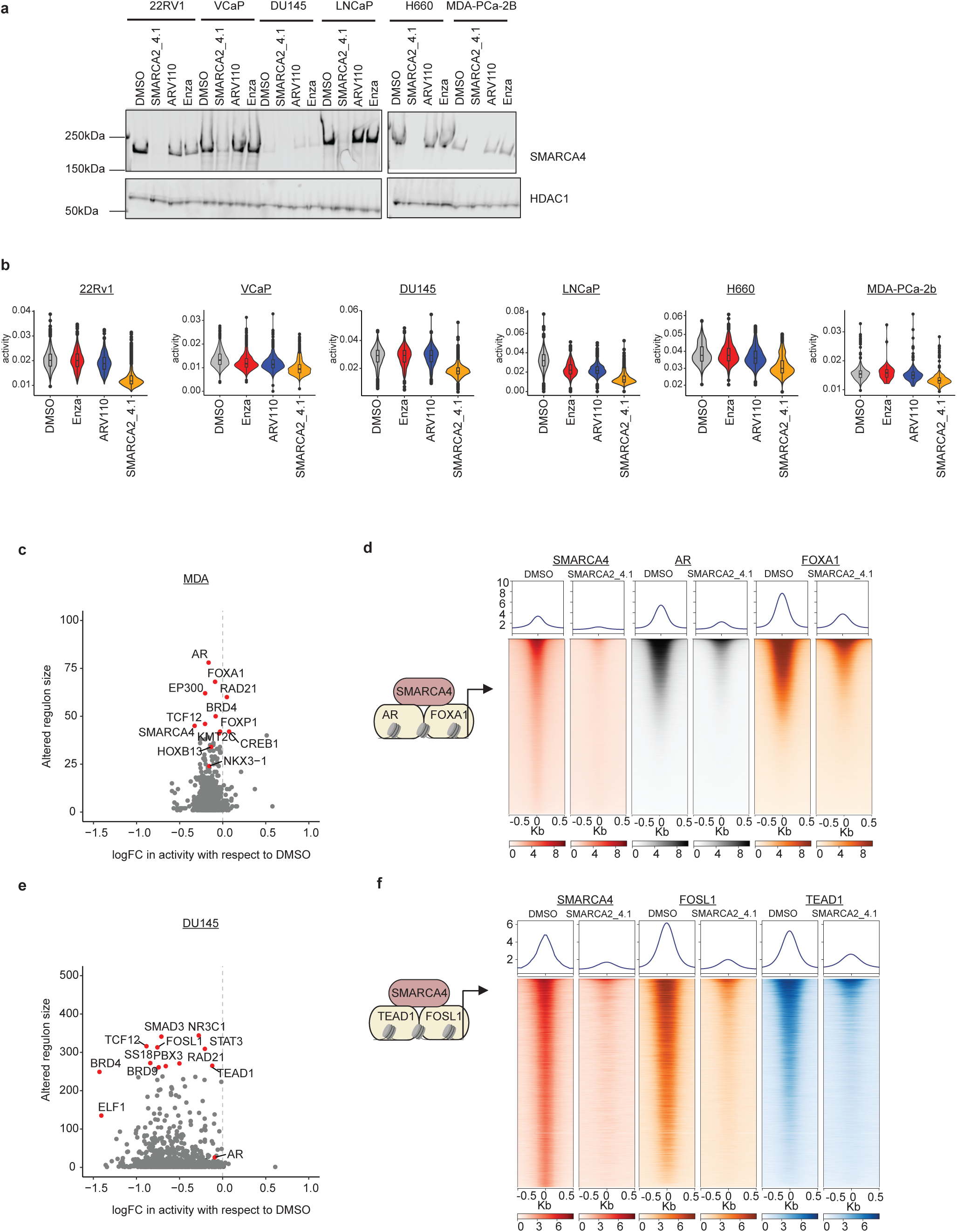
Epiregulon uncovers context-dependent interaction partners of SMARCA4. a) Immunoblotting of SMARCA4 after 24 hours of treatment and HDAC as a loading control b) SMARCA4 activity computed by Epiregulon for all 6 prostate cell lines after 24 hours of treatment c) Altered regulon size indicates the number of altered target genes mapped to each regulator. Altered genes are defined by genes with logFC > 0.5 and FDR <0.05 after SMARCA2_4.1 treatment. LogFC in activity indicates the changes in regulator activity estimated by Epiregulon. d) ChIP-seq of AR, SMARCA4 and FOXA1 in MDA-PCa-2b treated with the SMARCA2/4 degrader, SMARCA2_4.1 for 24 hours at 0.1 uM at regulatory elements mapped to SMARCA4 in the pruned regulon. e) Same as c) but for DU145 cells f) ChIP-seq of SMARCA4, TEAD1 and FOSL1 in DU145 cells treated with the SMARCA2/4 degrader, SMARCA2_4.1 for 24 hours at 0.1 uM

Even though the same starting SMARCA4 ChIP-seq data was used for all cell lines, Epiregulon constructed a different GRN for each cell line to capture the context-dependent effects of SMARCA2_4.1 treatment. In MDA-PCa-2b (an AR-dependent cell line), AR and FOXA1 were amongst the factors with the largest altered regulon size and differential activity (**Figure 4c**). ChIP-seq experiments indicated that SMARCA4 loss led to concomitant eviction of AR and FOXA1 at SMARCA4 binding sites (**Figure 4d**). In the AR-independent cell line DU145, FOSL1 and TEAD1 were amongst the top perturbed factors while AR was not (**Figure 4e**). ChIP-seq further validated the loss of FOLS1 and TEAD1 at SMARCA4 binding sites following SMARCA4 degradation (**Figure 4f**). This is consistent with their role in shaping the epigenomic landscape of stem-cell-like prostate cancer^19^. These results demonstrate how Epiregulon’s GRN can be inspected to identify context-specific cofactors of the targeted factor.

### Epiregulon identifies drivers of lineage reprogramming

To evaluate Epiregulon’s ability to identify drivers of cell states, we performed Reprogram-Seq^21^ to model lineage transition of prostate adenocarcinomas by overexpressing defined factors. Lentivirus encoding transcription factors were introduced into LNCaP cells using two independent constructs: 1) pLenti9-reprogram-seq-V2-Cbh-UTR2-3 (UTR) driven by the chicken beta-actin promoter containing a puromycin-resistant cassette, and 2) pLenti9-reprogram-seq-V2 (V2) driven by EF-1alpha promoter (**Figure 5a**). We overexpressed 4 factors, NKX2-1, GATA6, FOXA1 and FOXA2, along with a negative control mNeonGreen. NKX2-1 is a known driver of AR-independence and neuroendocrine transition in prostate cancer^22,23^ whereas GATA6 has unknown function in the prostate. FOXA2 promotes neuroendocrine transition in a genetically engineered mouse model^23^. None of these factors are expressed in parental LNCaP. FOXA1 is already expressed in parental LNCaP but we still introduced it since it is required for neuroendocrine transition^22^. We verified the expression of these factors using flow cytometry and immunoblotting. With the exception of NKX2-1 V2, other constructs achieved sufficient TF expression in LNCaP, with 30% to 70% of cells demonstrating overexpression above an uninfected control measured by flow cytometry (**Figure S5a-c**).

**Figure 5:**
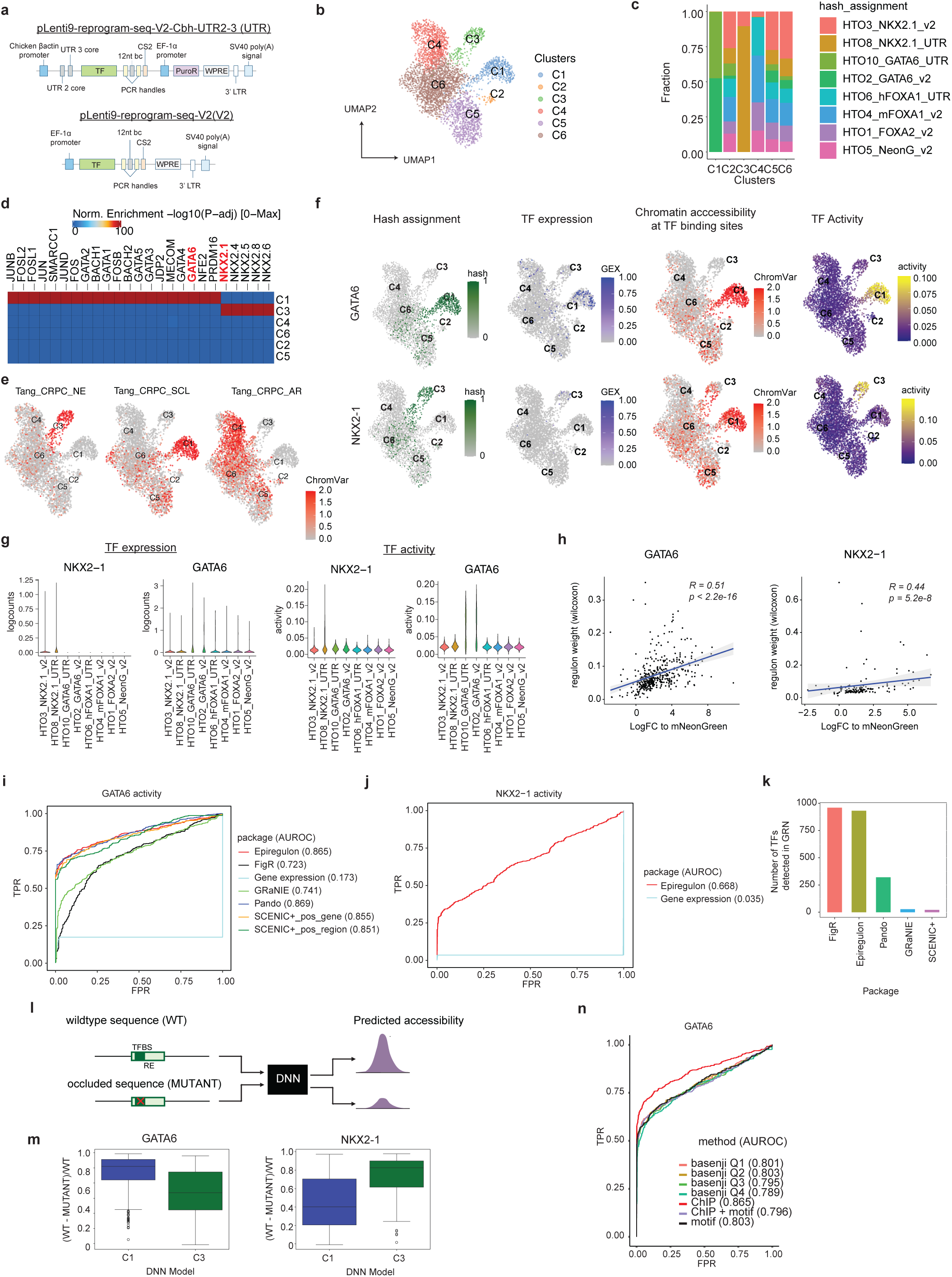
Epiregulon identifies drivers of lineage reprogramming. a) Two lentiviral constructs were used to introduce TF into LNCaP cells in the reprogram-seq assay. In pLenti9-reprogram-seq-V2-Cbh-UTR2-3, expression of TF is driven by the chicken beta-actin promoter and can be enriched with puromycin selection. pLenti9-reprogram-seq-V2, expression of TF is driven by EF-1alpha promoter. b) UMAP representation of 3,903 LNCaP cells transduced with virus encoding GATA6, NKX2-1, FOXA1, FOXA2 and mNeonGreen. The cells were infected in individual wells, hashtagged with HTO and then pooled into a single run. c) Distribution of HTO tags in each of the clusters. d) Motif enrichment in cluster specific peaks was performed by ArchR using the CisBP motif database. e) Chromatin accessibility at neuroendocrine (NE) - and stem cell like (SCL)-specific regions defined by Tang et al was computed by ChromVAR. f) Shown are the distribution of HTO tag assignment for GATA6 and NKX2-1, the level of TF gene expression, chromatin accessibility at GATA6-and NKX2-1 binding sites estimated by ChromVAR and TF activity computed by Epiregulon. g) Gene expression and Epiregulon-inferred activity of NKX2-1 and GATA6. h) Spearman correlation of regulon weights vs log fold changes of putative target genes of GATA6 (left) or NKX2-1 (right) with respect to mNeonGreen-infected cells i) Each cell was identified by the HTO tag corresponding to the well receiving virus encoding GATA6 or mNeonGreen and this information serves as the ground truth. A cell was classified into either expressing GATA6 or not based on its TF activity. AUROC was computed for all the packages being benchmarked as well as for GATA6 gene expression. Note that the theoretical maximum of AUROC will not reach 1 because not all cells took up virus encoding GATA6. j) Same as i) but for NKX2-1. Only Epiregulon could estimate the activity of NXK2-1. k) Number of TFs detected in the GRN computed by the different packages l) We train a deep neural network (DNN) model on the ATAC-seq signals from cluster 1 and cluster 3 cells respectively. We compare the predicted accessibility of either the wildtype sequence or the occluded sequence in the regulatory elements from the regulons inferred by Epiregulon. m) Normalized changes in predicted accessibility if we occlude the sequences found in the regulatory elements of GATA6 and NKX2-1 regulons in either the DNN model trained on cluster 1 or cluster 3 cells. n) Each cell was identified by the HTO tag corresponding to the well receiving virus encoding GATA6 or mNeonGreen and this information served as the ground truth cell labels. For each TF, a cell was classified into either expressing GATA6 or not. GATA6 ChIP-seq was obtained by merging ChIP-seq from ChIP-atlas and ENCODE. ChIP + motif refers to ChIP-seq peaks that contain GATA6 motifs. We trained a DNN model on cells expressing GATA6 (cluster 1) using Basenji2 and predicted an importance score for each motif based on the difference between the original sequence and motif occluded sequence. We filtered for those motifs with importance scores higher than quartiles of scores.

Expression of these TFs altered the cell state of LNCaP cells, leading to the formation of distinct clusters (**Figure 5b**). Cluster 1 was composed exclusively of GATA6-expressing cells while Cluster 3 contained only NKX2-1-expressing cells (**Figure 5c**). Furthermore, peaks upregulated in Cluster 1 and 3 were highly enriched for GATA6 and NKX2-1 motifs respectively (**Figure 5d**). NXK2-1 and GATA6 overexpression resulted in profoundly different cluster distributions from mNeonGreen (**Figure S5d**). Most strikingly, overexpression of NKX2-1 and GATA6 increased chromatin accessibility at neuroendocrine and stem-cell-like specific regions, respectively, and decreased accessibility at AR-dependent regions (**Figure 5e, S5e**) without overt changes in cell fitness (**Figure S5f**). We focused the rest of our analysis on GATA6 and NKX2-1 since their overexpression resulted in distinct reprogramming effects.

We tested Epiregulon’s ability to quantify the activity of GATA6 and NKX2-1 in this dataset. This is an interesting use case as the long distance between the polyA tail and the transcription factor cassette (1054bp in UTR and 2110bp in v2) precludes efficient capture of exogenous TF mRNA by the 3’ protocol used in scRNA-seq. As a result, the observed expression is a poor representation of TF activity (**Figure 5f, 5g**). In contrast, Epiregulon uses the expression of inferred target genes, correctly predicting increased activity for GATA6 in cluster 1 and NKX2 -1 in cluster 3 (**Figure 5f, 5g**). A subset of GATA6 targets were exclusively expressed in cluster 1 and NKX2-1 targets were exclusively expressed in cluster 3 (**Figure S5g**). In a differential expression analysis comparing cells in cluster 1 or 3 versus mNeonGreen control, GATA6 and NKX2-1 targets were highly ranked in cluster 1 and cluster 3, respectively (**Figure S5h**). Weights of target genes were also significantly correlated with their log fold changes (**Figure 5h**).

We performed a systematic benchmarking exercise to evaluate the performance of Epiregulon. All GRN methods can distinguish GATA6-expressing cells from mNeonGreen-expressing cells with similar accuracy (**Figure 5i**). However, Epiregulon was the only method that was able to predict NKX2-1 activity (**Figure 5j**). This improved sensitivity is attributed partially to the use of ChIP-seq data instead of motif annotations; the GRN derived from ChIP-seq data yielded 119 NKX2-1 targets, whereas no targets were obtained by Epiregulon with motif annotations. Indeed, Epiregulon’s GRN detected a large number of TFs mapped to their putative target genes (**Figure 5k**), highlighting the benefit of using ChIP-seq to empirically determine binding sites.

We used deep neural networks (DNNs) to determine the importance of sequence information within Epiregulon’s GRN. DNNs are capable of learning complex patterns in the input data associated with accessible regions to make predictions of ATAC-seq coverage^24–26^. We trained two DNN models with ATAC-seq signals from cluster 1 and cluster 3 cells respectively. We then occluded sequences within the REs in the regulons inferred by Epiregulon and compared the changes in predicted accessibility (**Figure 5l**). Occlusion of GATA6 REs resulted in greater alterations of chromatin accessibility in cluster 1 than cluster 3, while the converse was true for NKX2-1 (**Figure 5m**). This suggests that the REs inferred by Epiregulon contain important sequence information for determining accessibility.

We also considered whether the motif annotation approach could be improved by deep learning models. For GATA6, we computed motif importance scores with two independent sequence deep learning models, Basenji and ChromBPNet (see Methods. We could not perform this analysis in NKX2-1 due to the lack of REs passing significance based on motif annotations). We applied thresholds on these scores to identify the motifs that are most likely to correspond to binding sites. However, the use of thresholded motifs did not improve Epiregulon’s estimation of TF activity; in fact, overly stringent thresholding reduced the number of target genes and degraded performance (**Figure 5n** and **Figure S5i**). This motivates the continued use of the pan-cell-type ChIP-seq data, which still provides the best predictions of TF occupancy for activity inference.

### Epiregulon predicts known and novel drivers of the cancer state from clinical samples

We further applied Epiregulon to clinical specimens to evaluate its ability to discover regulators in heterogenous and complex samples. We obtained scATAC-seq and scRNA-seq data from primary tumors and normal adjacent tissues^27^ and used Epiregulon to construct a GRN on both normal epithelial and tumor cells for 3 different cancer indications (renal cell carcinoma, glioblastoma and pancreatic adenocarcinoma). Epiregulon detected many well-known factors including ZHX2, PAX8, N3RC1 in renal cell carcinoma, FOSL2 and SOX2 in glioblastoma and KLF5 in pancreatic adenocarcinoma (**Figure 6a-c** and refer to **Supplementary Table S4** for a full list of references). We also found that KLF9 was suppressed in pancreatic adenocarcinoma, consistent with its role in tumor suppression^28^. Interestingly, the changes in TF activity are more pronounced than changes in gene expression (**Figure 6a-c**). This implies that Epiregulon is able to identify drivers of the tumorigenic state even in the absence of strong changes in gene expression.

**Figure 6:**
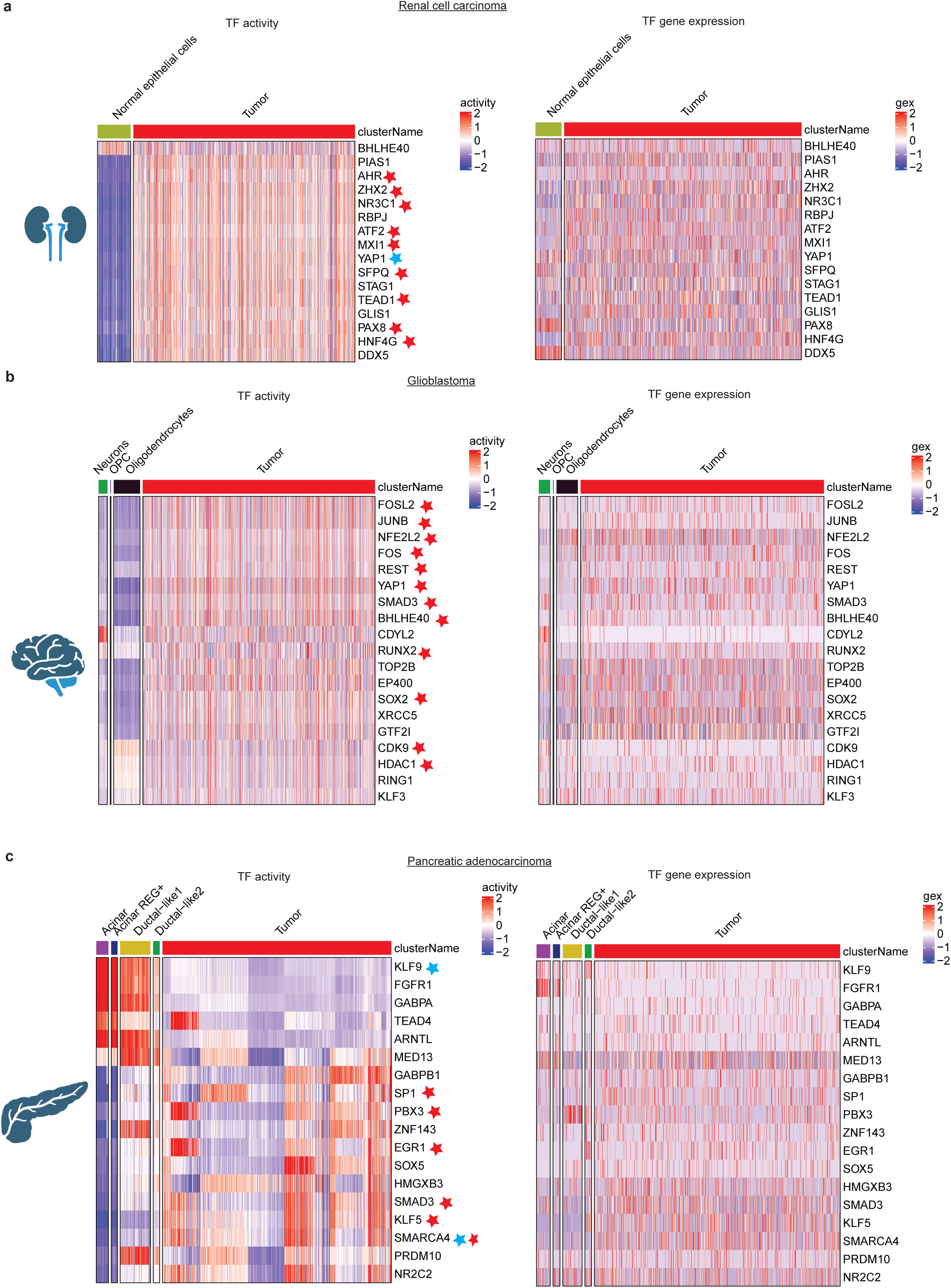
Epiregulon predicts known and novel drivers of the cancer state from clinical samples. ScATAC-seq and scRNA-seq data of primary tumors and normal adjacent tissues were obtained from Terekhanova et al^27^ and paired using Seurat’s label transfer function. Only tumor cells and matching normal cell types were used for GRN construction by Epiregulon (co-occurrence weight method). The top regulators were identified using Epiregulon’s *findDifferentialActivity* function. Shown are the expression and activity of the top regulators in a) renal cell carcinoma, b) glioblastoma and c) pancreatic adenocarcinoma. Red stars mark regulators which are known to promoter tumor growth and blue stars mark regulators which are known to inhibit tumor growth.

## Discussion

We present Epiregulon, a computational method to construct GRNs from single-cell multiomics data in a motif-agnostic manner. Epiregulon performs robustly across a multitude of datasets, identifying target genes and accurately quantifying TF activity even in the presence of neomorphic mutations. It is the most accurate tool for predicting response to drug perturbations and is the only tool that can infer the activity of a chromatin remodeler. We show that Epiregulon predictions based on ChIP-seq data outperform those from motif annotations, even after prioritization of the latter by DNN models. Our analyses demonstrate that Epiregulon can be reliably applied to evaluate TF-targeting pharmaceutical agents or to study epigenomic drivers of tumorigenesis and cell state changes.

Recent advances in chemical biology offer exciting new targeting strategies for the previously undruggable TFs and transcriptional coregulators. It will be important to develop a unified method to evaluate and compare various drug modalities which may include direct modulators of effector domain, degraders of protein and indirect modulators^29^. However, we have shown that TF expression is not a reliable measure of TF activity as negative feedback loops can still induce gene upregulation despite compromised TF function. Canonical gene signatures or motif-dependent gene regulatory network (GRN) methods also fail to account for alterations in the TF cistrome caused by gain-of-function mutations or TF hijacking ^30^ ^31^ ^32^. By constructing disease- or/and lineage-specific GRNs, Epiregulon helps identify the most likely targets for a particular model system and determines whether drugs are on-target and sufficiently potent. Furthermore, as demonstrated with the SMARCA4 degrader, Epiregulon can uncover therapeutically impactful co-targets. This GRN method can be applied to a broad range of motif-independent transcriptional coregulators including chromatin readers, transcriptional kinases and histone modifying enzymes, where identifying interaction partners is often challenging.

Epiregulon has some important limitations that may affect its performance. Acute perturbations can alter gene expression without substantial changes in chromatin accessibility^33^, reducing the effectiveness of Epiregulon and other GRN methods. Epigegulon does not explicitly model cooperativity between different TFs, which simplifies GRN construction but may reduce the accuracy of the activity estimates. Epiregulon also relies on the availability of ChIP-seq data, and while the pan-cell-type list is often satisfactory, best results are achieved by performing ChIP-seq on each TF of interest in the relevant biological system. Finally, our benchmarking was limited to the few TFs for which we have ground truth data, so it is difficult to generalize conclusions about Epiregulon’s performance to all TFs. Nevertheless, we envision that Epiregulon will become a useful tool for drug discovery, cancer biology and beyond.

## Methods

### Epiregulon

The Epiregulon workflow consists of several components: GRN construction, pruning of networks (optional), estimation of weights, calculation of activity, functional annotation of regulons, differential network analysis and identification of interaction partners. Epiregulon is designed to work seamlessly with ArchR but can accept any input as long as they are formatted into SingleCellExperiment objects. It is available as three related R packages. *Epiregulon* is the core package that performs GRN construction and activity inference. *Epiregulon.extra* contains differential analysis and plotting functions. *Epiregulon.archr* contains functions interfacing with ArchR. ChIP-seq data is available through *scMultiome*. All packages except *Epiregulon.archr* are available through Bioconductor.

#### Data preprocessing

Epiregulon assumes that prior preprocessing of the data has been performed by users’ methods of choice, requiring paired gene expression matrix (scRNA) and peak matrix (scATAC) along with dimensionality reduction matrix as the input. If gene expression and peak matrix are not paired, integration of scATAC-seq and scRNA-seq data must be performed prior to Epiregulon using methods such as ArchR’s addGeneIntegrationMatrix function.

For the datasets included in this manuscript, reads were mapped by the Cell Ranger ARC 2.0 and were further processed with the use of ArchR^34^. Briefly, cells were filtered based on TSS enrichment (>3) and number of ATAC-seq reads mapped to the nuclear genome (>1000). Moreover, doublets were removed using ArchR’s filterDoublets function. ATAC-seq data was represented as a 500bp tile matrix on which iterative latent sematic indexing (LSI) was performed. Similarly, normalized gene expression data was subject to iterative LSI. The dimensionality reductions from both modalities were then combined into a single matrix that was used for cell clustering by graph clustering approach implemented in Seurat^35–37^. The ATAC-seq peaks were called separately for each cluster and after normalization were merged to produce one peak set for all cells (peak x cell matrix). The cells were assigned to their sample barcodes using demuxEM^38^. Enrichment of chromatin accessibility at each peak or motif was computed using the chromVAR^39^ function provided through ArchR. Briefly, chromVAR computes the bias-corrected deviation of per-cell accessibility for a given motif or TF binding sites from the average of all cells.

#### Network construction

Epiregulon provides two similar methods to establish peak to gene links using the calculateP2G function. If there is an existing ArchR project, Epiregulon retrieves the peak to gene links that have been previously assigned by ArchR. Briefly, ArchR creates 500 cell aggregates, resampling cells if needed. ArchR computes the correlation between chromatin accessibility and target genes within a size window (default ±250 kb) and retains peak-gene pairs that exceed a correlation threshold (default Epiregulon cutoff is 0.5). In the absence of an ArchR project, Epiregulon defines cell aggregates using k-means clustering based on the reduced dimensionality matrix and performs correlation in the same manner as ArchR does. If cluster labels are provided, cluster - specific correlations are reported in addition to overall correlations.

#### TF occupancy data

Each regulatory element is then interrogated for TF occupancy based on a compilation of public TF ChIP-seq binding sites. ChIP-seq were downloaded from ChIP-Atlas (chip-atlas.org) and ENCODE (encodeproject.org). Only ChIP-seq peaks with FDR < 1 x 10^-5^ were retained. We created a pan-cell-type ChIP-seq peakset using the merged ChIP-seq peaks provided by ChIP-Atlas and the ENCODE ChIP-seq data, yielding 1557 unique factors (human) and 768 unique factors (mouse). We also created sample- and tissue-specific ChIP-seq peak sets for each factor to allow for tissue or sample matched analysis. We only retained ChIP-seq samples that meet the following criteria:

- Total number of unique reads >= 10M
- Percentage mapped reads >= 70%
- Number of peaks (FDR< 1 x 10^-5^) >= 100

Peak sets were merged across each sample or each tissue for every TF. Data is provided in the scMultiome package as GRanges list objects and can be accessed using the getTFMotifInfo function from the Epiregulon package or directly from the scMultiome package using the tfBinding function.

If desired, ChIP-seq peaks can be further annotated for the presence of motifs using the addMotifScore function. If motif annotation has been performed previously using ArchR’s addMotifAnnotations function, motif annotations can be easily retrieved and appended to the peak matrix. In the ArchR independent workflow, Epiregulon can annotate peak matrix using motifmatchr’s motif matching function^39^ and cisbp as the reference motif database^40^. Alternatively, users can start entirely from motif annotations and leverage deep learning motifs to select motifs based on motif importance scores (see section on DNN models).

#### Network pruning (optional)

Epiregulon prunes the network by performing tests of independence on the observed number of cells jointly expressing transcription factor (TF), regulatory element (RE) and target gene (TG) vs the expected number of cells if TF/RE and TG are independently expressed using the pruneRegulon function.

We define 𝑛 as the total number of cells, 𝑘 as the number of cells jointly expressing TF, TG and RE above a set threshold, 𝑔 as the number of cells jointly expressing TF and RE above a threshold and ℎ as the number of cells expressing TG above a threshold. 𝑝, the expected probability of cells jointly expressing TF, TF and RE above a threshold is defined as follows:

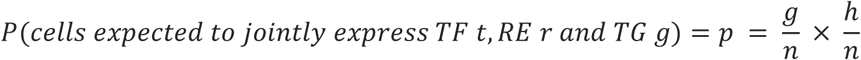

Two tests of independence are implemented, the binomial test and the chi-square test. In the binomial test, the expected probability is 𝑝, the number of trials is the total number of cells 𝑛, and the observed number of successes is 𝑘, the number of cells jointly expressing all three elements.

In the chi-square test, the expected probability for having all 3 elements active is also 𝑝. The observed cell count for the active category is 𝑘, and the cell count for the inactive category is 𝑛 −𝑘. P-values are calculated from chi-squared distribution with degree of freedom equal to 1. Cluster-specific p-values are calculated if users supply cluster labels. Finally, multiple hypothesis testing was performed by the Holm method.

#### Estimation of weights

While network pruning provides statistics on the joint occurrence of TF-RE-TG, we would like to further estimate the strength of regulation using the addWeights function. Biologically, this can be interpreted as the magnitude of gene expression changes induced by changes in transcription factor activity. Epiregulon provides 3 different methods to estimate weights. Two measures (correlation and co-occurrence (Wilcox)) give both the magnitude and directionality of changes whereas weights computed by mutational information (MI) are always non-negative. Within 2 of the methods (correlation and MI), there is an option of modeling the TG expression based only on TF expression, or on the product of TF expression and RE chromatin accessibility. Consideration of both TF expression and RE chromatin accessibility is highly recommended, especially for scenarios in which TF activity and TF gene expression are decoupled (as in the case of drug perturbations or CRISPR genome editing). The correlation and mutual information statistics are computed on pseudobulks by user-provided cluster labels and yield a single weight across all clusters per each TF-RE-TG triplet. In contrast, the Wilcoxon method groups cells based on the joint expression of TF, RE and TG in each single cell or in cell aggregates. Cell aggregation uses a default value of 10 cells and can help overcome sparsity and speed up computation. If cluster labels are provided, we can obtain cluster-specific weights using the Wilcoxon method.

##### 1) Co-occurrence (Wilcox)

Cells were divided into two groups, with the first group jointly expressing TF gene expression and chromatin accessibility at RE, and the remaining cells in the second group. The default cutoff is 1 for normalized gene expression and 0 for normalized chromatin accessibility. There is also an option to use the median of each feature as an adaptive cutoff.

##### 2) Correlation

If we consider only TF expression (tf_re.merge set to FALSE), the weight is the correlation coefficient between the TF gene expression and TG gene expression. If we consider both TF expression and RE chromatin accessibility (tf_re.merge set to TRUE), the weight is the correlation coefficient between the product of TF gene expression and RE chromatin accessibility vs the TG gene expression.

##### 3) Mutual information between the TF and target gene expression

If we consider only TF expression (tf.re_merge is set to FALSE), the weight is the mutual information between the TF gene expression and the TG gene expression. If we consider both TF expression and RE chromatin accessibility (tf.re_merge is set to TRUE), the weight is the mutual information between the product of TF gene expression and RE chromatin accessibility vs the TG gene expression.

#### Calculation of TF activity

The activities for a specific TF in each cell are computed by averaging the weighted expressions of target genes linked to the TF:

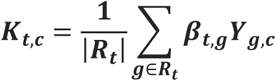

where 𝐾_𝑡,𝑐_ is the activity of a TF 𝑡 for a cell 𝑐, |𝑅_𝑟_| is the total number of target genes for a TF 𝑡, 𝑌_𝑔_ is the normalized count of target gene 𝑔 where 𝑔 is regulated by TF 𝑡 and 𝛽_𝑡,𝑔_ is the regulatory weight of TF 𝑡 on target gene 𝑔. 𝑅_𝑡_ is the regulon of TF 𝑡. If cluster labels are provided, cluster-specific weights are used.

#### Gene set enrichment of regulons

Gene set enrichment of a regulon is performed by testing whether the target genes of a TF are over-represented in known gene signatures such as those provided by MSigDB using a hypergeometric test. Target genes can be refined by filtering the regulons on user-defined weights.

#### Differential TF activity by total activity

This differential analysis compares the differences in the activity of each transcription factor between conditions. This analysis is well suited for identifying factors that have contrasting levels of activities, for instance, lineage factors that are turned on or off during certain developmental stages or cell types. TF activities are compared between groups using any standard statistical methods. Here we use scran’s findMarkers function to find differential activity between user provided groups of cells (https://rdrr.io/bioc/scran/man/findMarkers.html).

#### Differential TF activity by network topology

A second approach to investigate differential TF activity is to compare and contrast target genes or network topology. This is useful when a transcription factor differs in the target genes it regulates, while maintaining a similar level of total activity. This can happen when a transcription factor redistributes to a different set of genomic regions, due to mutations in the transcription factors or changes in the interaction partners. Differences in network topology are calculated by taking the degree centrality of the edge subtracted graphs between two conditions, with cluster-specific regulon weights representing edge weights in each condition.

Consider networks 𝐺(1) and 𝐺(2) with identical node sets N and respective adjacency matrices **A**(1) and **A**(2). Then then the edge-subtracted network 𝐺′ induced by 𝐺(1) and 𝐺^(2)^ has the adjacency matrix whose entries are calculated as 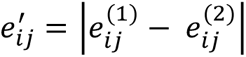, where 𝑒^(𝑗)𝑖𝑗^ represents the weight of edge connecting *i*th transcription factor with *j*th gene in network 𝐺^(𝑗).^

We simplify the tripartite TF-RE-TG graph to a bipartite TF-TG graph by taking the maximum of TF-RE-TG weights of the same TF-TG pairs. Degree centrality is the sum of the weights associated with the *i*th transcription factor: 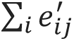. Degree centrality is further normalized to account for differences in the number of target genes of each transcription factor. The default normalization method is dividing degree centrality by the square root of the number of target genes. This strikes a balance between penalizing TFs with an abundance of target genes and prioritizing TFs with differential target genes. Transcription factors are ranked by normalized degree centrality.

### Benchmarking

#### Benchmarking using PBMC data

PBMC dataset was downloaded from 10x genomics (https://www.10xgenomics.com/datasets/pbmc-from-a-healthy-donor-granulocytes-removed-through-cell-sorting-10-k-1-standard-2-0-0). The Cell Ranger output was processed using ArchR as described in the data preprocessing section. As a result, 9,702 out of the 11,582 cells were kept for downstream analysis. We used peak matrix retrieved from ArchR project as input to GRN inference tools. Gene expression data was retrieved from ArchR or directly from Cell Ranger output, depending on the benchmarked tool.

Clustering was performed using LSI dimensionality reduction, which combined information from both chromatin accessibility and gene expression data. We used marker genes to determine naive CD4+ T cells (IL7R, CCR7), CD14+ monocytes (CD14, LYZ), and CD4+ memory cells (S100A4, IL7R) and SingleR with BlueprintEncodeData from celldex package as a reference for other cell types. Cell clusters were annotated into one of the following types: naive CD4+ T, memory CD4+ T, naive CD8+ T, memory CD8+ T, monocytes, CD14+ monocytes, FCGR3A+ monocytes, B, NK, DC. 24 cells were left unannotated and were excluded from further analyses.

We used PBMC data from the KnockTF database^41^ as the ground truth to test accuracy of target genes assignment to transcription factors. We collected data presenting the results of seven knock-down experiments, each one targeting a different transcription factor. Target genes were determined using quality filters (absolute value of logFC > 0.5, corrected p-value < 0.05). For each package we benchmarked, we calculated precision and recall based on the predicted and ground truth target genes. Precision is defined as the number of predicted target genes that are altered by the knockdown of TF / total number of predicted target genes. Recall is defined as the number of predicted target genes that are altered by the knockdown of TF / total number of altered genes.

All packages were tested for run time and memory use. Data preprocessing was excluded from these measurements (including topic analysis in SCENIC+). We assigned 64GB and 20 cores on HPC node for each run. In the case of GRaNIE the memory allocation had to be increased to 128 GB and for FigR the memory allocation was increased to 256 GB. Each package was run 5 times and median run time and memory use were recorded.

#### Benchmarking using Reprogram-Seq data

We introduced 4 transcription factors into LNCaP cells - FOXA1, FOXA2, NKX2-1 and GATA6 and obtained paired gene expression and chromatin accessibility information. Because FOXA1 was already highly expressed in LNCaP cells at the basal level, its introduction did not have any profound impact on lineage plasticity and therefore FOXA1 was excluded in subsequent analyses. We focused on NKX2-1 and GATA6 because their expression resulted in distinct cell clusters. Each cell is identified by the HTO tag corresponding to the well receiving virus encoding a particular TF and this information serves as the ground truth cell labels. For each TF, a cell is classified into either expressing this TF or not. AUROC was computed for all the packages being benchmarked as well as for gene expression of the TF being evaluated. Note that because each TF has different transduction efficacy and not all cells are able to take up the virus or express the TF, the theoretical maximum of AUROC will not reach 1.

#### Benchmarking using AR antagonist data

For the drug treatment dataset, we treated 6 prostate cancer cell lines with 3 different therapeutic agents and obtained paired gene expression and chromatin accessibility information. Only the 2 AR-dependent cell lines, LNCaP and VCaP, were included in the benchmark since they are known to respond to AR-targeting agents. Similarly, only enzalutamide^1^ and ARV-110^3^ were used in the benchmark studies since they are known to specifically inhibit AR activity. Cells treated with Enzalutamide and ARV110 are supposed to show reduced AR activity compared to cells treated with DMSO control. Each cell line was analyzed separately by retaining only peaks found in each cell line. Each cell is identified by the HTO tag corresponding to the well receiving DMSO or one of the two AR inhibitors and this information serves as the ground truth. For each of the two drugs (enzalutamide or ARV110), a cell is classified into either treated with DMSO control or an AR inhibitor. AUROC is computed for all the packages being benchmarked, based on the AR activity values, as well as for AR gene expression. In the Epiregulon workflow we used the Wilcoxon method to estimate weights with cell-line AR ChIP-seq.

We used 4 GRN inference tools (FigR, SCENIC+, *GRaNIE* and *Pando)* to benchmark the performance of Epiregulon. To ensure consistency, we applied the same gene expression and chromatin accessibility matrices across all tools. We first used ArchR to determine the peaks i.e. DNA regions with frequent Tn5 transposase insertion events from the fragment files output by CellRanger. Then the peak x cell matrix was produced by counting the number of insertions per peak per cell. From the same ArchR project we also retrieved normalized gene expression to be used by Epiregulon, GRaNIE, FigR. For the remaining tools, we used a subset of cells in the gene expression data which matched the cells in the ArchR project to make sure that all tools work on the same cells. Below we describe the workflows used in each tool. We followed the tutorial examples provided at official web sites with only minor changes to the default settings.

#### FigR

We performed benchmarking against FigR^9^. In the first step of GRN construction the correlation between peak accessibility and target gene expression was determined using runGenePeakcorr. We used a non-default search range around TSS (250 kb) to be consistent across all the benchmarked tools. Correlation coefficients were also computed with background peaks and the significance of gene-peak association was determined with one-tailed z-test. Only gene-peak associations that show positive correlation and are statistically significant were retained (p ≤ 0.05). DORaCs (High density domains of regulatory chromatin) correspond to genes with ≥7 associated peaks and were calculated by summing the ATAC-seq reads in peaks matched to each gene. Each TF is associated with a single motif by selecting the motifs most highly correlated with other motifs of the same TF. Each DORC is evaluated for enrichment of TFs by comparing the match frequency in its peakset vs the match frequency in a set of background peaks; the p-values of the enrichment were obtained by z-test (𝑃_𝐸𝑛𝑟𝑖𝑐ℎ𝑚𝑒𝑛𝑡_). Smoothing of gene expression matrix and chromatin accessibility data summarized across target genes was performed after determination of k nearest neighbor cells based on LSI space retrieved from ArchR project and constructed using ATAC-seq data. The Spearman correlations between the smoothed DORC accessibility and smoothed TF gene expression were computed (𝐶𝑜𝑟𝑟𝑒𝑙𝑎𝑡𝑖𝑜𝑛) and their significance were obtained using z-test (𝑃_𝐶𝑜𝑟𝑟𝑒𝑙𝑎𝑡𝑖𝑜𝑛_ ). From the output of the main function (getFigRGRN) we retrieved scores indicating the strength of association between transcription factors and target genes which were computed as below:

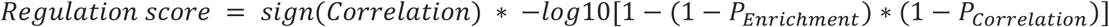

The regulation scores were used as weights when calculating activity with Epiregulon.

#### SCENIC+

We performed benchmarking against SCENIC+^11^. Briefly, we used pycisTopic which uses Latent Dirichlet Allocation to group regulatory elements into the topics. The model evaluation indicated 20, 20, 10, 10 as the number of topics for VCaP, LNCaP, MDA and 22Rv1 cells from AR dataset, respectively and 20 for Reprogram-Seq dataset. The input peak matrix was retrieved from the AchR project. We then used pycistarget to identify TF-region links. Pycistarget identifies motif matches in the peak regions using HOMER, scores each region for motif importance and identifies differentially enriched motifs above background regions. Only regions showing NES > 3.0 and motifs with adjusted p-value <0.05 and log2FC >0.5 were retained. TF-gene importance scores were calculated using gradient-boosting machine regression by predicting TF expression from target gene expression. Region-gene importance scores were calculated using gradient-boosting machine regression by predicting target gene expression from region accessibility. Genes were ranked by TF-gene importance scores and only genes in the leading edge of the gene set enrichment were used for the eRegulon. Gene set enrichment was also performed for region-gene pairs. Peaks were ranked by imputed chromatin accessibility and genes were ranked by gene expression counts per cell. Enrichment score was defined as the AUC at 5% of the ranking and was calculated using AUCell. The enrichment score was used as the activity score in the benchmark assessment.

#### GRaNIE

We performed benchmarking against GRaNIE^10^. Briefly, GRaNIE overlapped TF binding sites obtained from HOCOMOCO-based TF motifs with ATAC-seq peaks. GRaNIE identified TF-peak connections using Pearson correlation between TF expression and the peak accessibility across samples. The cutoffs for the correlation were chosen based on an empirical FDR calculated from the ratio of TF-peaks in the background peaks over the total number of TF-peaks in both the background and foreground. Peak-gene connections were identified using correlation between the gene expression and chromatin accessibility. All the GRN edges have a weight of 1. The default threshold is FDR < 0.2 for TF-peak links and FDR < 0.1 for peak-gene links. Activities were computed using Epiregulon’s calculateActivity.

#### Pando

We performed benchmarking against Pando^8^. Briefly, ATAC-seq peaks were intersected with PhastCons conserved elements and cCREs derived from ENCODE. TFs present in the 4000 most variable genes were included in the downstream analysis. Motifs for the TFs were obtained from JASPAR2020 and CISBP. Gene expression and chromatin accessibility counts were smoothened by averaging cells within a neighborhood. Pando then models target gene expression as the weighted sum of the product of TF expression and the chromatin accessibility of the region where the TF binds to.

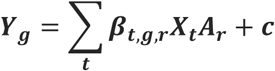

where 𝑌_𝑔_ is the expression of the target gene 𝑔, 𝑋_𝑡_ is the expression of the transcription factor 𝑡, 𝐴_𝑟_ is the chromatin accessibility at region 𝑟, 𝛽 is the fitted coefficient and 𝑐 is the intersection.

Fitted coefficients were tested for significance using analysis of variance (ANOVA). Only edges with FDR <0.05 were retained in the final GRN. Activities were calculated using Epiregulon’s calculateActivity function with the fitted coefficients 𝛽_𝑡,𝑔,𝑟_ as the weights.

### GRN construction from patient tumors

Unpaired scATAC-seq fragment files and Seurat objects containing author-processed scRNA-seq counts of primary tumors and normal adjacent tissues were downloaded from NCI Human Tumor Atlas Network as indicated in Terekhanova et al^27^. scATAC-seq data was preprocessed by ArchR as described in the data preprocessing section. Pairing of scATAC-seq and scRNA-seq was performed using Seurat’s label transfer function implemented within ArchR with patient ID as the restraint. Only tumor cells and matching normal cell types were retained and were down sampled to 50,000 cells for each indication. GRN was constructed using Epiregulon’s co-occurrence (wilcox) weight estimation method and the top regulators were identified using Epiregulon’s *findDifferentialActivity* function. Only TFs with altered regulon size greater than 35 were retained. Altered regulon size refers to the number of genes showing an absolute normal-tumor log fold change > 0.5 and FDR <0.05.

### DNN training

Steps for DNN-based TF-RE mapping involved dataset processing, model training, motif scoring and thresholded GRN construction.

#### Dataset processing

The reference genome hg38 and the pseudo-bulk ATAC (taken from ArchR) of individual clusters (cluster 1 or cluster 3) was collected. The whole genome excluding unmappable regions^43^ was split into 3072 bp regions. Using the coordinates of each region the corresponding DNA sequence and bigwig coverage was obtained. The DNA sequence was converted to one-hot encoded form. The DNA and coverage information were saved in tf records files for faster I/O during training. Training, validation and test splits were done by chromosomes taking chr8 as test set and chr9 as validation set.

#### Model training

We used Basenji2 architecture with 32 base-pair resolution (binned coverage at 32 bp). The model was trained by randomly sampling 2048bp segment from the input DNA and taking the corresponding coverage (one cluster per model). We used Poisson NLL as loss, used reverse complement augmentation and trained for a maximum of 50 epochs with early stopping.

We additionally trained models with base-pair resolution using the ChromBPNet architecture. We first trained a custom Tn5 sequence bias model for this dataset on non-peak regions. We ensured that these models learned Tn5 bias by applying DeepSHAP to the model outputs. Using this bias model to regress out Tn5 sequence bias, we then trained ChromBPNet TF models. These models were trained on 2114bp sequences centered on ATAC-seq peaks.

#### Motif scoring

For each of the motif regions (taken from ArchR motif positions file) we obtained the DNA sequence (denoted as wild type) by centering at the motif and extending to 2048bp. We occluded the motif by replacing motif nucleotides with ‘N’ (or 0.25 in one-hot encoding) (denoted as mutant). We computed the score - mean fraction change in predicted coverage by subtracting mutant from wild type and normalizing by predicted wild type coverage.

#### Entire RE occlusion

Similarly, we assessed the importance of entire REs for model comparison. We used ChIP-based regulons to identify genomic coordinates containing GATA6 (or NKX2-1) binding sites, at 500bp resolution. We then obtained wild-type (WT) predictions for sequences centered at these REs and mutant sequence predictions by occluding the entire 500bp RE. We quantified the importance of each RE by subtracting the sum of the mutant predictions from the WT predictions and normalizing by the WT value.

#### Thresholded GRN construction

We computed several quartiles of scores as thresholds. We filtered for those motifs with importance scores higher than the threshold corresponding to bigger differences in prediction. Using the RE remaining we constructed the GRN as before.

### Reprogram-Seq

#### Construction of Lentivirus Plasmids for TF Over-expression

All lentivirus plasmids were generated by GenScript. Briefly, the ORF of transcriptional factors were codon optimized, synthesized and cloned after the hEF1a promoter. The puromycin resistance gene is driven by a separate Cbh promoter to enable antibiotics selection. Maxi-prep of each plasmid was performed to maximize the transfection efficiency.

#### Cell Culture and Virus Packaging

LNCaP Clone FGC cells were obtained from ATCC and cultured with RPMI-1640 media with 10% FBS and 2mM L-Glutamine. The cells were split every 4-5 days to maintain the appropriate density. 293T cells were cultured in DMEM with 10% FBS, 100μM NEAA, 2mM Glutamine. They were split every 2-3 days. One day before transfection, the cell culture dish was treated with 5mL 1% gelatin in PBS, incubated for 10 minutes, then aspirated. 3.5X10^6^ 293T cells were seeded into each 10 cm dish to reach ∼80% confluence. On the day of transfection, 20 µl Lipofectamine 2000 was added to 480 µl OptiMEM. In a new tube, the plasmid was mixed in 500µl Opti-MEM at the following ratio: carrier plasmid, 5 µg; delta8.9, 16 µg; VSVG, 1 µg. Both mixes were combined and incubated for 20 minutes before adding to the dish. The dish was incubated at 37°C for 6 hours and then 6 ml of complete media was added. Virus was then harvested 48 hours after transfection. Briefly, all supernatants were harvested and filtered through a 0.45 µm filter bottle. The virus was concentrated by using the Lenti-X concentrator (TAKARA Bio.) following the manufacturer’s instructions and resuspended in 1 ml 1% BSA in PBS per dish. The concentrated virus was divided into 200µl aliquots and stored in -80°C until infection.

#### Infection of LNCaP Cells by Lentivirus

Two days before infection, 4X10^5^ LNCaP cells were seeded into each well of a 6-well plate. On the day of infection, aspirate the media and change to 0.5mL RPMI + 10% FBS + 1X Glutamax with 8 µg/ml Polybrene. A total of 200 µl concentrated lentivirus was added to each well to achieve high MOI, and then the plate was centrifuged at 1800 rpm for 45 minutes at room temperature. After that the plate was put into the incubator for another 3 hours before 2 ml of the full media was added to each well. Two days after infection, the media was refreshed with 1µg / ml puromycin. The cells were then grown for another 7 days before harvesting for the single-cell analysis, during which the cells were split accordingly when the confluence reached 100%.

#### Engineering of dox-inducible NKX2-1 expressing cell lines

LNCAP Clone FGC cells were passaged two days before electroporation. 200k cells per reaction were transduced with 375 ng PBIND-NKX2-1-IRES-eGFP-PGK-puro and 125ng pBO (PiggyBac transposase) (3:1 vector:transposase ratio) according to manufacturer’s instructions for SF Cell Line 4D-Nucleofector™ X Kit S, using the EN-120 electroporation protocol in 20 μl Nucleocuvette™ Strip. After electroporation, 100ul of warm media (RPMI + 10% FBS + 1X Glutamax) was added to the cuvette and the cuvette was placed in the incubator for 1 hour to recover. Cells were then transferred into a 6 well plate. Puromycin selection (RPMI + 10% FBS + 1X Glutamax, 1ug/ml Puromycin) was performed after 4-5 days, depending on cell recovery.

### Drug treatment with AR inhibitors and SMARCA2/4 degrader

LNCap Clone FGC, VCAP and DU145 cells were obtained from ATCC and cultured with RPMI-1640 media with 10% FBS and 2mM L-Glutamine; NCI-H660 cells were obtained from ATCC and cultured with DMEM/F12 with 0.005 mg/ml insulin, 0.01 mg/ml Transferrin, 30nM Sodium selenite,10nM Hydrocortisone,10nM beta-estradiol, 4mM L-Glutamine. The cells were split every 4-5 days to maintain the appropriate density. For the drug response assay, cells were seeded in 6 well format. The cells were split every 4-5 days to maintain the appropriate density.

1.5x10^5^ DU145 cells were seeded into each well of a 6 well plate and 5E3 DU145 cells were seeded into each well of a 96 well plate; 1.5-2E6 LNCAP or VCAP cells were seeded into T75 flasks; 5x10^5^ cells NCI-H660 cells were seeded into each well of a 6 well plate. After 48 hours, the media was removed and cells were treated with DMSO, 1uM Enzalutamide, 0.1 uM SMARCA2_4.1 or 1 uM ARV-110 in complete media. After 24 hours from treatment, the cells were harvested for the single-cell RNA-ATAC Co-assay

### Single-cell RNA-seq and Single-cell RNA-ATAC Co-assay

The single-cell RNA-seq was performed using the Chromium Single Cell 3’ kit (V3.1) from 10X Genomics with cell hashing. Briefly, the cells were trypsinized into single-cell suspension from the 6-well plates, and washed once with PBS with 1% BSA. The cells from different wells were stained with human TotalSeq-A cell hashing antibodies (BioLegend) containing distinct barcodes following the manufacturer’s protocol. The cells were then washed twice with PBS with 1% BSA before combining and the live cells were sorted using the SONY SH800S FACS machine. For loading of the 10X chip G, we overloaded the channel with the aim of recovering 20K cells. The library construction was following the standard 10X protocol with the following exceptions: 1) the 1µl of 10µM HTO additive primer was spiked in during the cDNA PCR step; 2) during the SPRI beads cleanup step, the supernatant from the 0.6X cleanup was saved to recover the HTO fragment. Along with the transcriptome library, the HTO library was amplified from the supernatant using the HTO-specific primer. The detailed protocol can be found on the cell hashing website (https://cite-seq.com/cell-hashing/).

The single-cell RNA-ATAC co-assay was performed using the 10X Genomics Single Cell Multiome ATAC + Gene Expression kit. In a previous test, we found that the standard cell permeabilization protocol used for nuclei preparation does not completely remove the plasma membrane of the LNCap cells and robust B2M expression can still be detected with FACS (data not shown). Therefore, we speculated that the same cell hashing strategy would work for the 10X Multiome assay. Briefly, we followed the same hashing protocol as used in the scRNA-seq experiment. After the antibody staining and pooling, the cells were permeabilized by following the nuclei isolation protocol from 10X genomics ([10X Nuclei Isolation Protocol](https://www.10xgenomics.com/support/single-cell-multiome-atac-plus-gene-expression/documentation/steps/sample-prep/nuclei-isolation-for-single-cell-multiome-atac-plus-gene-expression-sequencing)). The cells were then counted and loaded on the Chip J with a recovery aim of 12K cells per lane. For the library construction, we followed the standard protocol with the following modifications: 1) 1µl of 10µM HTO additive primer was spiked in during the pre-AMP and cDNA PCR steps; 2) during the 0.6X cDNA cleanup, the supernatant was saved to amplify the HTO library, as described in the previous section.

All the libraries were sequenced on Illumina NextSeq 2000 or NovaSeq 4000. We were aiming for 20K, 40K and 2K raw reads per cell for the transcriptome, ATAC and HTO libraries, respectively.

### Synthesis of SMARCA2_4.1

**Scheme 1.**
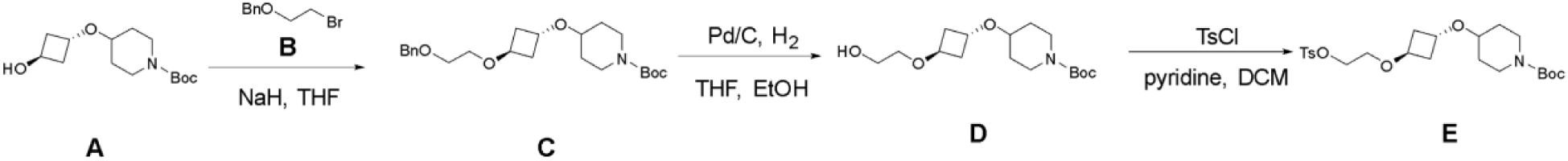
Preparation of intermediate E.

*tert-Butyl 4-((trans)-3-(2-(benzyloxy)ethoxy)cyclobutoxy)piperidine-1-carboxylate. (**C**)*. To a solution of *tert*-butyl 4-((trans)-3-hydroxycyclobutoxy)piperidine-1-carboxylate (**A**, 500 mg, 1.84 mmol, 1.0 eq) in THF (7 mL) was added NaH (60% in mineral oil, 111 mg, 2.77 mmol, 1.5 eq) at 0 °C. The resulting solution was stirred at 0 °C for 0.5 h afterwhich ((2-bromoethoxy)methyl)benzene (**B**, 592 mg, 2.76 mmol, 1.5 eq) was added. The mixture was warmed to room temperature and was stirred for an additional 16 h then was quenched with saturated aqueous NH_4_Cl (5 mL). The resulting solution was extracted with EtOAc (10 mL x 3), and the combined organic layers were dried over sodium sulfate and concentrated under vacuum. The residue was purified by silica gel chromatography (EtOAc:petroleum ether, 0:100 to 24:76) to afford the title compound (330 mg, 44%) as a colorless oil. ^1^H NMR (300 MHz, DMSO-*d*_6_) δ 7.42-7.23 (m, 5H), 4.49 (s, 2H), 4.25-4.17 (m, 1H), 4.09-4.02 (m, 1H), 3.68-3.61 (m, 2H), 3.57-3.51 (m, 2H), 3.47-3.37 (m, 2H), 2.95 (t, *J* = 11.0 Hz, 2H), 2.23-1.98 (m, 4H), 1.79-1.67 (m, 2H), 1.39 (s, 9H), 1.35-1.17 (m, 3H). LC-MS: (ES+): m/z 406 [M+H]^+^.

*tert-Butyl 4-((trans)-3-(2-hydroxyethoxy)cyclobutoxy)piperidine-1-carboxylate. (**D**)*. A solution of intermediate **C** (330 mg, 0.815 mmol) and 10% Pd/C (66.0 mg) in ethanol (10 mL) and THF (10 mL) was stirred at room temperature for 1 h under a hydrogen atmosphere (3 atm). The resulting mixture was filtered, and the filter cake was washed with DCM (10 mL x 2). The filtrate was concentrated under reduced pressure to afford the title compound (245 mg, 95% yield) as a white oil. ^1^H NMR (300 MHz, DMSO-*d*_6_) δ 4.58 (t, *J* = 5.5 Hz, 1H), 4.26-4.18 (m, 1H), 4.08-4.01 (m, 1H), 3.69-3.61 (m, 2H), 3.53-3.34 (m, 3H), 3.28 (t, *J* = 5.3 Hz, 2H), 2.96 (t, *J* = 11.4 Hz, 2H), 2.24-1.98 (m, 4H), 1.76-1.71 (m, 2H), 1.39 (s, 9H), 1.36-1.18 (m, 2H). LC-MS: (ES+): m/z 316 [M+H]^+^.

*tert-Butyl 4-((1r,3r)-3-(2-(tosyloxy)ethoxy)cyclobutoxy)piperidine-1-carboxylate (**E**)*. To a solution of intermediate **D** (245 mg, 0.78 mmol, 1.0 eq) and pyridine (245 mg, 3.11 mmol, 4.0 eq) in DCM (2 mL) was added TsCl (223 mg, 1.16 mmol, 1.5 eq) at 0 °C. The mixture was then warmed to room temperature and stirred for 12 h. The solvent was subsequently removed under vacuum, and the residue was purified by silica gel chromatography (EtOAc:petroleum ether, 0:100 to 29:71) to afford the title compound (303 mg, 83%) as a colorless oil. ^1^H NMR (300 MHz, DMSO-*d*_6_) δ 7.82-7.72 (m, 2H), 7.52-7.41 (m, 2H), 4.22-4.05 (m, 3H), 4.01-3.88 (m, 1H), 3.67-3.59 (m, 2H), 3.45-3.34 (m, 3H), 2.94 (t, *J* = 11.4 Hz, 2H), 2.41 (s, 3H), 2.13-1.92 (m, 4H), 1.76-1.64 (m, 2H), 1.37 (s, 9H), 1.33-1.15 (m, 2H). LC-MS: (ES+): m/z 470 [M+H]^+^.

**Scheme 2.**
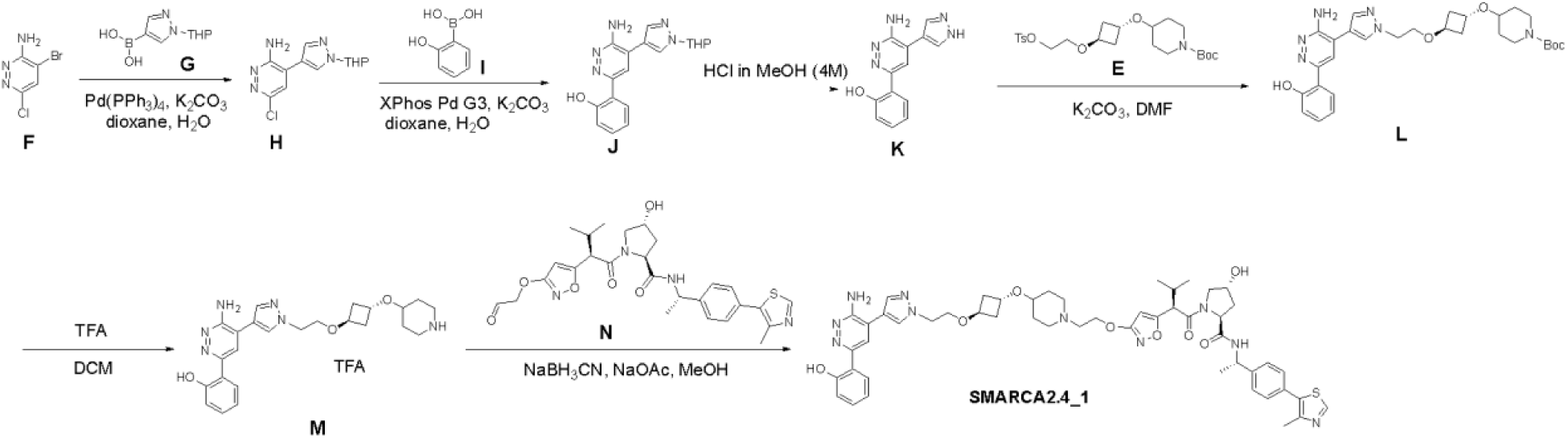
Preparation of SMARCA2_4.1 (compound 1).

*6-Chloro-4-(1-(tetrahydro-2H-pyran-2-yl)-1H-pyrazol-4-yl)pyridazin-3-amine (**H**)*. A solution of 4-bromo-6-chloropyridazin-3-amine (**F**, 2.00 g, 9.60 mmol, 1.0 eq), (1-(tetrahydro-2H-pyran-2-yl)-1H-pyrazol-4-yl)boronic acid (**G**, 2.46 g, 12.5 mmol, 1.3 eq), Pd(PPh3)4 (1.12 g, 0.969 mmol, 0.1 eq) and K_2_CO_3_ (2.67 g, 19.3 mmol, 2.0 eq) in 1,4-dioxane (20 mL) and water (2 mL) was stirred at 100 °C for 2 h. The reaction was then cooled to room temperature and was quenched with water (20 mL). The resulting solution was extracted with EtOAc (30 mL x 3), and the combined organic layers were dried over sodium sulfate and concentrated under vacuum. The residue was purified by silica gel chromatography (EtOAc:petroleum ether, 0:100 to 58:42) to afford the title compound (2.40 g, 89%) as a red oil. ^1^H NMR (300 MHz, DMSO-*d*_6_) δ 8.52 (d, *J* = 0.8 Hz, 1H), 8.11 (d, *J* = 0.8 Hz, 1H), 7.65 (s, 1H), 6.41 (s, 2H), 5.46 (dd, *J* = 9.9, 2.2 Hz, 1H), 4.07-3.90 (m, 1H), 3.76-3.57 (m, 1H), 2.24-2.05 (m, 1H), 2.00-1.91 (m, 2H), 1.79-1.64 (m, 1H), 1.60-1.53 (m, 2H). LC-MS: (ES+): m/z 280 [M+H]^+^.

*2-(6-Amino-5-(1-(tetrahydro-2H-pyran-2-yl)-1H-pyrazol-4-yl)pyridazin-3-yl)phenol (**J**)*. A solution of intermediate **H** (1.35 g, 4.83 mmol, 1.0 eq), (2-hydroxyphenyl)boronic acid (**I**, 870 mg, 6.31 mmol, 1.3 eq), XphosPdG3 (410 mg, 0.484 mmol, 0.1 eq) and K_2_CO_3_ (1.33 g, 9.64 mmol, 2.0 eq) in 1,4-dioxane (11.5 mL) and water (2.5 mL) was stirred at 95 °C for 5 h. The mixture was cooled to room temperature, and the solvent was removed under vacuum. The residue was purified by silica gel chromatography (MeOH:DCM, 0:100 to 5:95) to afford the title compound (900 mg, 55%) as a yellow solid. ^1^H NMR (300 MHz, DMSO-*d*_6_) δ 13.77 (s, 1H), 8.60 (d, *J* = 0.8 Hz, 1H), 8.26-8.17 (m, 2H), 8.02 (dd, *J* = 8.3, 1.6 Hz, 1H), 7.28-7.23 (m, 1H), 6.97-6.85 (m, 2H), 6.51 (s, 2H), 5.48 (dd, *J* = 9.9, 2.2 Hz, 1H), 3.99-3.90 (m, 1H), 3.74-3.60 (m, 1H), 2.25-2.06 (m, 1H), 2.00-1.95 (m, 2H), 1.76-1.64 (m, 1H), 1.59-1.52 (m, 2H). LC-MS: (ES+): m/z 338 [M+H]^+^.

*2-(6-Amino-5-(1H-pyrazol-4-yl)pyridazin-3-yl)phenol (**K**)*. A solution of intermediate **J** (300 mg, 0.890 mmol) in HCl (4.0 M in MeOH, 1 mL) was stirred at room temperature for 2 h. The solvent was then removed under vacuum, and the residue was purified by pre-packed C18 column (solvent gradient: 0-100% MeOH in water (0.1% TFA)) to afford the title compound (188 mg, 83%) as a brown solid. ^1^H NMR (300 MHz, DMSO-*d*_6_) δ 13.85 (s, 1H), 13.42 (s, 1H), 8.33 (s, 2H), 8.22 (s, 1H), 8.02 (dd, *J* = 8.3, 1.6 Hz, 1H), 7.30-7.24 (m, 1H), 6.95-6.90 (m, 2H), 6.50 (s, 2H). LC-MS: (ES+): m/z 254 [M+H]^+^.

*tert-Butyl 4-((trans)-3-(2-(4-(3-amino-6-(2-hydroxyphenyl)pyridazin-4-yl)-1H-pyrazol-1-yl)ethoxy)cyclobutoxy)piperidine-1-carboxylate (**L**)*. A solution of intermediate **K** (100 mg, 0.395 mmol, 1 eq), intermediate **E** (223 mg, 0.475 mmol, 1.2 eq) and K_2_CO_3_ (136 mg, 0.985 mmol, 2.5 eq) in DMF (1.3 mL) was stirred at 80 °C for 2 h. The reaction mixture was cooled to room temperature, then was diluted with EtOAc (5 mL) and washed with brine (3 mL x 5). The organic layer was dried over sodium sulfate and was concentrated under vacuum. The residue was purified by silica gel chromatography (MeOH:DCM, 0:100 to 5:95) to afford the title compound (60 mg, 27%) as a brown solid. ^1^H NMR (300 MHz, DMSO-*d*_6_) δ 13.79 (s, 1H), 8.45 (d, *J* = 0.8 Hz, 1H), 8.24-8.17 (m, 2H), 8.03-7.97 (m, 1H), 7.25 (t, *J* = 7.7 Hz, 1H), 6.97-6.87 (m, 2H), 6.48 (s, 2H), 4.30 (t, *J* = 5.4 Hz, 2H), 4.15-4.06 (m, 1H), 4.06-4.00 (m, 1H), 3.69 (t, *J* = 5.4 Hz, 2H), 3.61-3.53 (m, 2H), 3.24-3.13 (m, 1H), 2.84 (t, *J* = 11.3 Hz, 2H), 2.05-2.01 (m, 4H), 1.66-1.60 (m, 2H), 1.35 (s, 9H), 1.24-1.17 (m, 2H). LC-MS: (ES+): m/z 551 [M+H]^+^. *2-(6-Amino-5-(1-(2-(trans)-3-(piperidin-4-yloxy)cyclobutoxy)ethyl)-1H-pyrazol-4-yl)pyridazin-3-yl)phenol, TFA salt (**M**)*. A solution of intermediate **L** (60 mg, 0.11 mmol) in TFA (0.3 mL) and DCM (0.3 mL) was stirred at room temperature for 0.5 h. The solvent was removed under vacuum to afford the title compound (66 mg, crude) as a yellow solid. ^1^H NMR (300 MHz, DMSO-*d*_6_) δ 8.46 (s, 1H), 8.26 (s, 1H), 8.14 (s, 1H), 7.71-7.68 (m, 1H), 7.39-7.33 (m, 1H), 7.03-6.91 (m, 2H), 4.32 (t, *J* = 5.4 Hz, 2H), 4.20-4.00 (m, 2H), 3.70 (t, *J* = 5.4 Hz, 2H), 3.53-3.45 (m, 1H), 3.12-3.08 (m, 2H), 2.90-2.84 (m, 2H), 2.09-2.05 (m, 4H), 1.91-1.78 (m, 2H), 1.59-1.47 (m, 2H). LC-MS: (ES+): m/z 451 [M+H]^+^.

*(2S,4R)-1-((R)-2-(3-(2-(4-((trans)-3-(2-(4-(3-amino-6-(2-hydroxyphenyl)pyridazin-4-yl)-1H-pyrazol-1-yl)ethoxy)cyclobutoxy)piperidin-1-yl)ethoxy)isoxazol-5-yl)-3-methylbutanoyl)-4-hydroxy-N-((S)-1-(4-(4-methylthiazol-5-yl)phenyl)ethyl)pyrrolidine-2-carboxamide (**1**)*. A solution of intermediate **M** (66 mg, crude), intermediate **N** (prepared as described in Berlin, M.; et al. “PROTACs Targeting BRM (SMARCA2) Afford Selective In Vivo Degradation Over BRG1 (SMARCA4) and are Active in BRG1-Mutant Xenograft Tumor Models”, *J. Med. Chem*. accepted for publication; 85 mg, 0.157 mmol) and NaOAc (54 mg, 0.658 mmol) in MeOH (0.8 mL) was stirred at room temperature for 1 h. Then NaBH_3_CN (21.0 mg, 0.33 mmol) was added, and the mixture was stirred at room temperature for an additional 0.5 h. The solvent was removed under vacuum, and the residue was purified by Prep-HPLC [column: Xselect CSH F-Phenyl OBD column, 30*150 mm, 5 μm; mobile Phase A: water (10 mmol/L NH_4_HCO_3_), mobile phase B: CH_3_CN; flow rate: 60 mL/min; gradient: 43% to 63% B over 8 min; wavelength: 254/220nm; RT: 8.97 min] to afford the title compound (12.8 mg, 12% over two steps) as a white solid. ^1^H NMR (300 MHz, DMSO-*d*_6_) δ 13.83 (s, 1H), 9.00-8.99 (m, 1H), 8.51-8.46 (m, 1H), 8.43 (d, *J* = 7.6 Hz, 1H), 8.26-8.19 (m, 2H), 8.05-8.01 (m, 1H), 7.47-7.34 (m, 4H), 7.29-7.24 (m, 1H), 6.98-6.87 (m, 2H), 6.50 (s, 2H), 6.09-5.92 (m, 1H), 5.11-5.02 (m, 1H), 4.96-4.86 (m, 1H), 4.44-4.24 (m, 4H), 4.15 (t, *J* = 5.6 Hz, 2H), 4.11-4.00 (m, 2H), 3.74-3.51 (m, 4H), 3.46-3.43 (m, 1H), 3.14-3.08 (m, 1H), 2.66-2.63 (m, 2H), 2.57-2.53 (m, 2H), 2.46 (s, 3H), 2.31-2.16 (m, 1H), 2.07-1.94 (m, 7H), 1.82-1.74 (m, 1H), 1.67-1.63 (m, 2H), 1.46-1.37 (m, 3H), 1.34-1.20 (m, 2H), 0.98 -0.95 (m, 3H), 0.84-0.79 (m, 3H). LC-MS: (ES+): m/z 975 [M+H]^+^.

## Acknowledgments

We would like to thank Xinpeng Wang, Fang Zhang, Mingtao He, Yunxing Cheng and Michael Berlin (Arvinas) for the synthesis of SMARCA2_4.1 and Peter Dragovich for guiding the synthesis. We would like to thank Jayaram Kancherla and Amber Schedlbauer for their help and advice.

## Supplementary Figures

**Figure S1**

RNA and ATAC counts of similar cells are first aggregated. Based on the correlation between ATAC and RNA, each target gene (TG) is assigned a regulatory element (RE) within a defined distance (default ±250kb). Each RE is interrogated for possible TF occupancy using ChIP-seq data from ENCODE and ChIP-Atlas. An optional network pruning step retains TF-RE-TG triplets that co-occur within the same cells. Co-occurrence is defined by TF and TG expression and chromatin accessibility exceeding a threshold. The regulatory strength of TF for each TG is estimated using 1 of the 3 methods, correlation, mutual information and co-occurrence (Wilcoxon) and this serves as the weight of each TF-TG pair. The weights are computed for all the cells and/or for cells in each cluster to yield cluster-specific weights. The TF activity is computed as the weighted sum of the target gene expression normalized by the number of genes with non-zero edges. The regulon of each TF can be annotated with gene set enrichment. Jaccard similarity between the target genes is used to identify transcription factors that potentially interact. Differential network is performed on edge-subtracted networks of two cell states.

**Figure S2**

Individual precision and recall of 7 factors as described in Figure 1f

**Figure S3**

a) Cell viability assessed by cellTiter-Glo at 24 hours after treatment
b) AUROC of the AR dataset was computed using AR activity as the classifier of treatment conditions: DMSO or AR inhibitor (ARV110 or enzalutamide). The two AR-dependent cell lines (VCaP and LNCaP) are known to decrease AR activity upon ARV110 or enzalutamide treatment.
c) The same as b) but shown for two resistant cell lines (22RV1 and MCa-PCa-2b). Both cell lines do not respond to enzalutamide and respond only mildly to ARV110.

**Figure S4**

a) Viability of MDA-PCa-2b cells stimulated by varying concentrations of hydrocortisone or 5α-DHT over 5 days.
b) AR activities in MDA-PCa-2b cells calculated from AR signature genes (Dong^16^, Hieronymus^17^, Nyquist^18^, Tang et al^19^)
c) Overlap of AR ChIP-seq peaks in VCaP versus MDA-PCa-2b
d) Overlap of putative AR targets in VCaP versus MDA-PCa-2b
e) The ground truth regulator activity was computed using all the publicly available ChIP-seq obtained in VCaP cells. This activity was then correlated (Pearson’s) with activity computed either from pan-cell-type ChIP-seq (red) or motif annotations (blue) for each of the regulators.
f) Correlation of activity obtained from pan-cell-type ChIP-seq vs activity obtained from cell line matched ChIP-seq. Each point represents a cell.
g) Shown is the Jaccard similarity between the target genes derived from LNCaP ChIP-seq vs target genes derived from pan-cell-type ChIP-seq (red) or motif annotations (blue).

**Figure S5**

a) Intracellular flow cytometry showing the expression of GATA6, NKX2-1 and FOXA2 driven by 2 independent constructs, pLenti9-reprogram-seq-V2-Cbh-UTR2-3 (UTR23) or pLenti9-reprogram-seq-V2 (V2) in LNCaP. Expression of mNeonGreen was also shown as a control.
b) Percentage of cells showing increased expression of the indicated factor compared to uninfected control
c) Immunoblotting of NKX2-1 and GATA6 in LNCaP cells. Both TFs were driven by pLenti9-reprogram-seq-V2-Cbh-UTR2-3. NCI-H660 was used as a positive control for GATA6 and NCI-H226 was used as a positive control for NKX2-1.
d) Distribution of cluster assignment in each treatment condition
e) Correlation of GATA6 activity versus chromatin accessibility at stem cell like (SCL)-specific regions present in castration resistant prostate cancer (CRPC) samples reported by Tang et al (left). Correlation of NKX2-1 activity versus chromatin accessibility at neuroendocrine (NE)-specific regions in CRPC tumors (right). Each points represents a single cell and is colored by its cluster assignment. Increased chromatin accessibility at SCL-specific regions is highly enriched for Cluster 1 cells and increased accessibility at NE-specific regions are enriched in Cluster 3 cells.
f) Growth rate of parental LNCaP cells and LNCaP cells overexpressing GATA6 or NKX2-1 measured by Incucyte.
g) Normalized expression of GATA6 and NKX2-1 putative target genes inferred by Epiregulon
h) Genes were ranked by a product of two ranks: one based on the differences in the mean expression between TF-expressing group and control groups, and the second based on the proportion of the total gene expression across all the cells contributed by the treated cells. Gene set enrichment was computed using the NKX2-1 target genes and GATA6 target genes respectively.
i) Same as Figure 5n but performed with chromBPNet

## Code availability

Epiregulon: https://github.com/xiaosaiyao/epiregulon

Epiregulon.extra: https://github.com/xiaosaiyao/epiregulon.extra

Epiregulon.archr: https://github.com/xiaosaiyao/epiregulon.archr

Epiregulon.book: https://xiaosaiyao.github.io/epiregulon.book/

ScMultiome: https://github.com/xiaosaiyao/scMultiome

Epiregulon manuscript: https://github.com/xiaosaiyao/epiregulon.manuscript

Deep neural network model: https://github.com/xiaosaiyao/epiregulon.sequence.modeling

Differential network simulation: https://github.com/xiaosaiyao/epiregulon.diffnetwork.simulation

## Data availability

GEO (GSE252883)

GSE251977:

https://www.ncbi.nlm.nih.gov/geo/query/acc.cgi?acc=GSE251977

token: snczwmievvwnbyl

GSE251978:

https://www.ncbi.nlm.nih.gov/geo/query/acc.cgi?acc=GSE251978

token: sdkpsuwutdoblep

