## Supplementary Figures for "Epiregulon: Inference of single-cell transcription factor activity to predict drug response and drivers of cell state"

Figure S1: Epiregulon constructs GRNs to infer TF activity at the single cell level

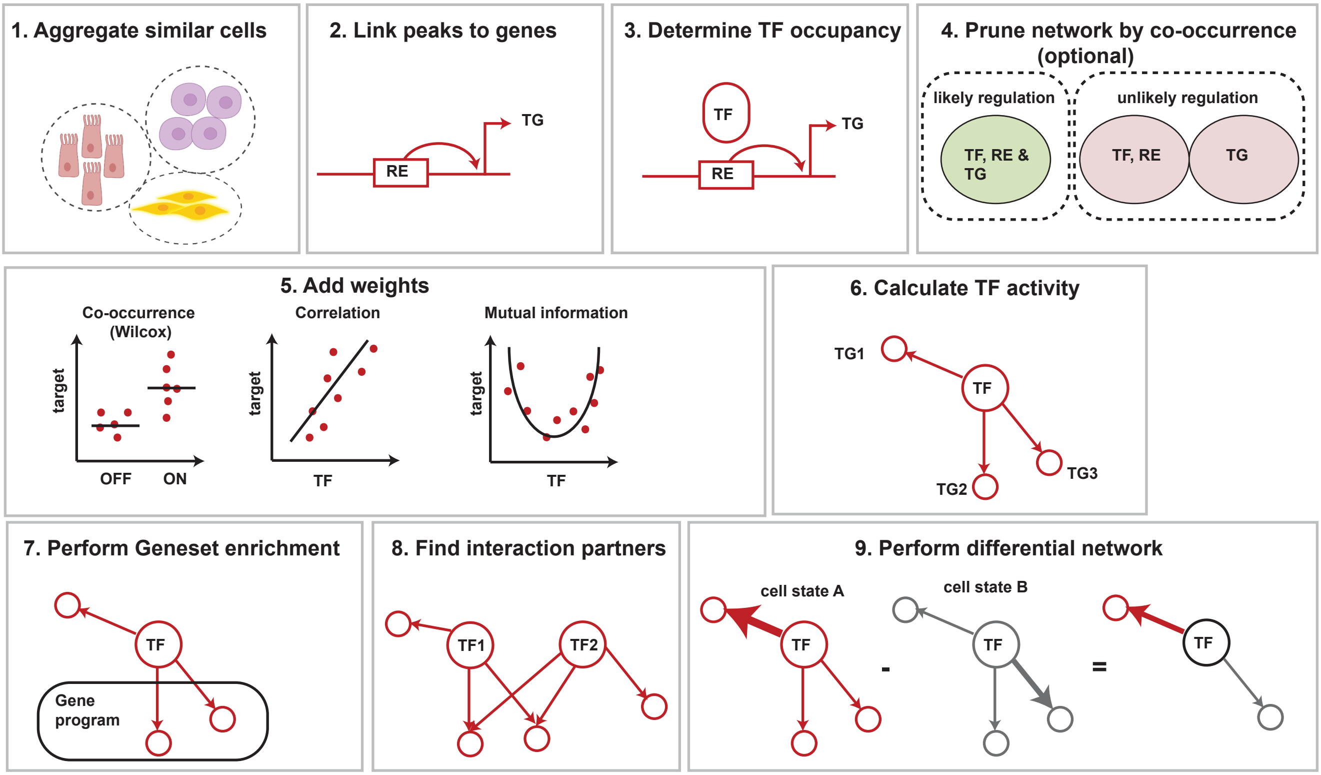

**Figure S2**

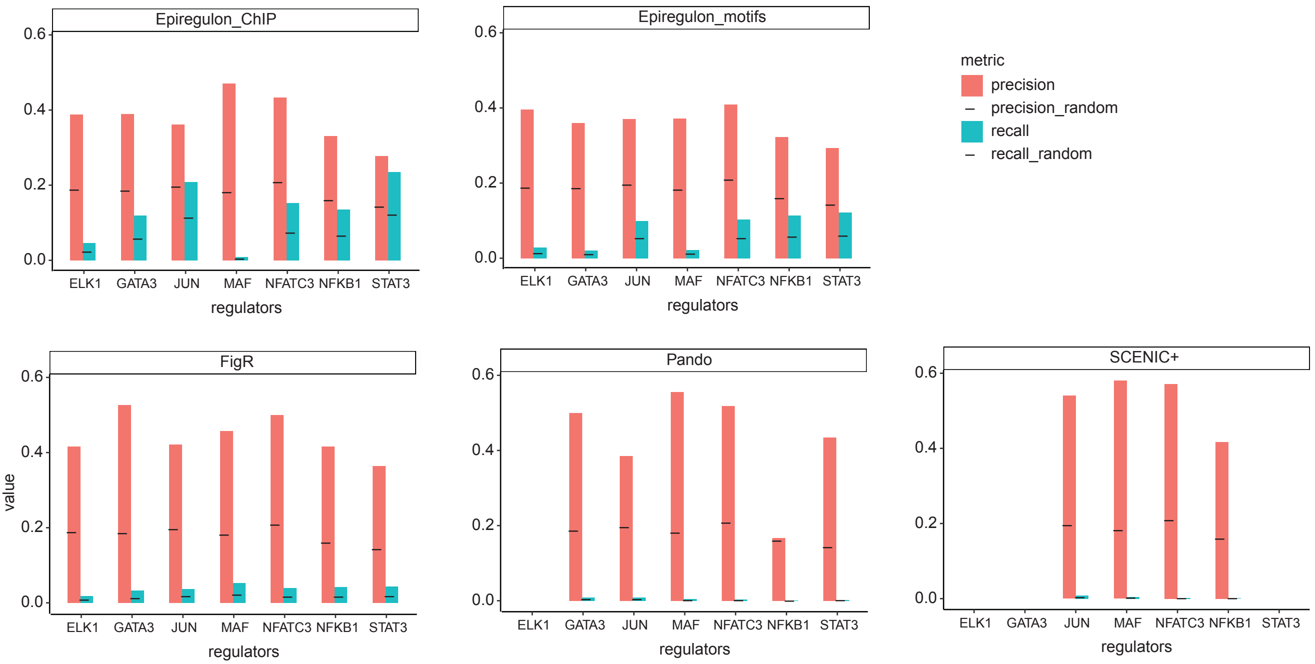

Epiregulon_ChIP
Epiregulon_motifs
FigR
Pando
SCENIC+
value
regulators
ELK1
GATA3
JUN
MAF
NFATC3
NFKB1
STAT3
0.0
0.2
0.4
0.6
metric
precision
precision_random
recall
recall_random

**Figure S3**

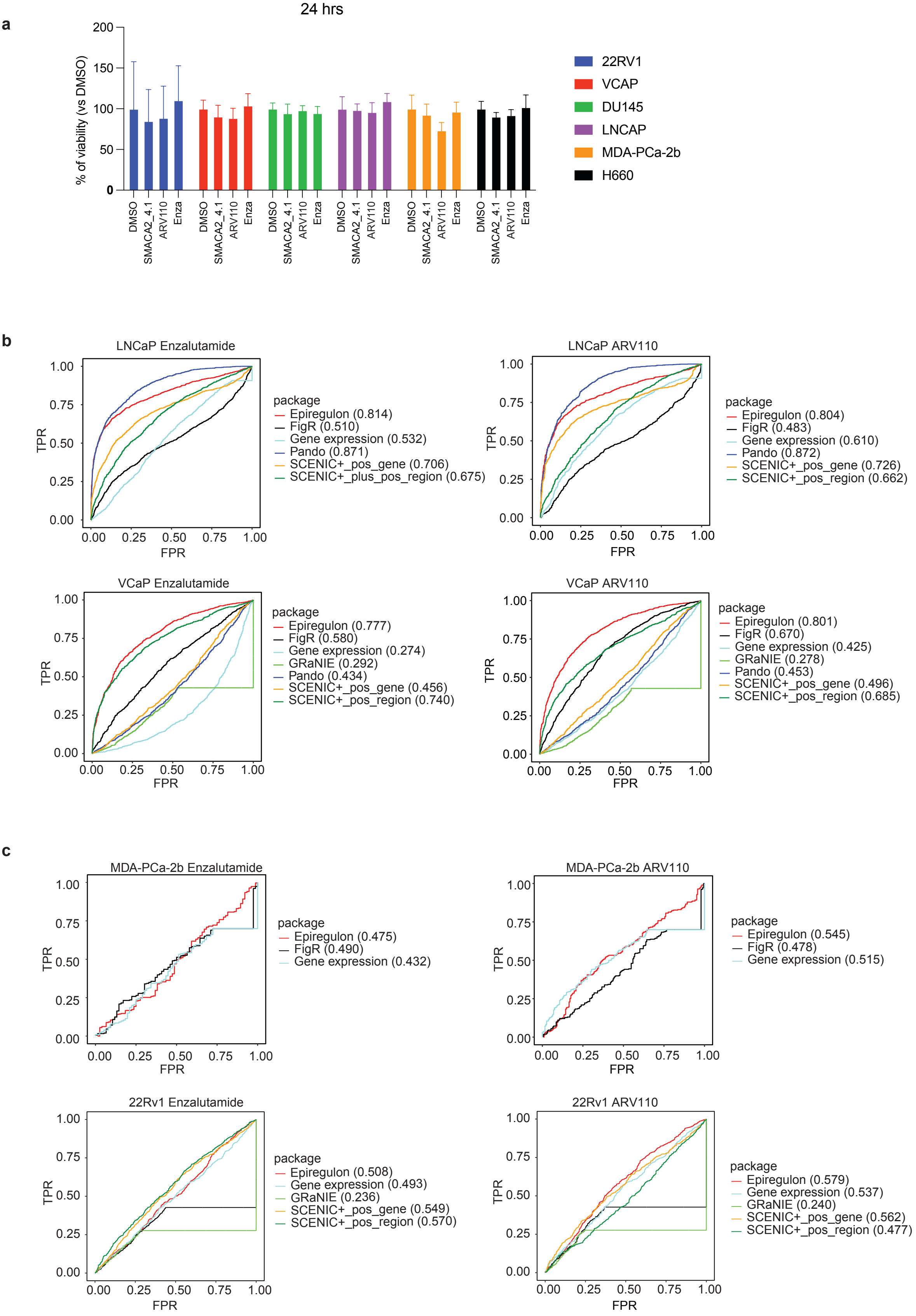
a
24 hrs
% of viability (vs DMSO)
200
150
100
50
0
DMSO
SMACA2_4.1
ARV110
Enza
22RV1
VCAP
DU145
LNCAP
MDA-PCa-2b
H660
b
LNCaP Enzalutamide
package
Epiregulon (0.814)
FigR (0.510)
Gene expression (0.532)
Pando (0.871)
SCENIC+_pos_gene (0.706)
SCENIC+_plus_pos_region (0.675)
LNCaP ARV110
package
Epiregulon (0.804)
FigR (0.483)
Gene expression (0.610)
Pando (0.872)
SCENIC+_pos_gene (0.726)
SCENIC+_pos_region (0.662)
VCaP Enzalutamide
package
Epiregulon (0.777)
FigR (0.580)
Gene expression (0.274)
GRaNIE (0.292)
Pando (0.434)
SCENIC+_pos_gene (0.456)
SCENIC+_pos_region (0.740)
VCaP ARV110
package
Epiregulon (0.801)
FigR (0.670)
Gene expression (0.425)
GRaNIE (0.278)
Pando (0.453)
SCENIC+_pos_gene (0.496)
SCENIC+_pos_region (0.685)
TPR
FPR
0.00
0.25
0.50
0.75
1.00
c
MDA-PCa-2b Enzalutamide
package
Epiregulon (0.475)
FigR (0.490)
Gene expression (0.432)
MDA-PCa-2b ARV110
package
Epiregulon (0.545)
FigR (0.478)
Gene expression (0.515)
22Rv1 Enzalutamide
package
Epiregulon (0.508)
Gene expression (0.493)
GRaNIE (0.236)
SCENIC+_pos_gene (0.549)
SCENIC+_pos_region (0.570)
22Rv1 ARV110
package
Epiregulon (0.579)
Gene expression (0.537)
GRaNIE (0.240)
SCENIC+_pos_gene (0.562)
SCENIC+_pos_region (0.477)

Figure S4

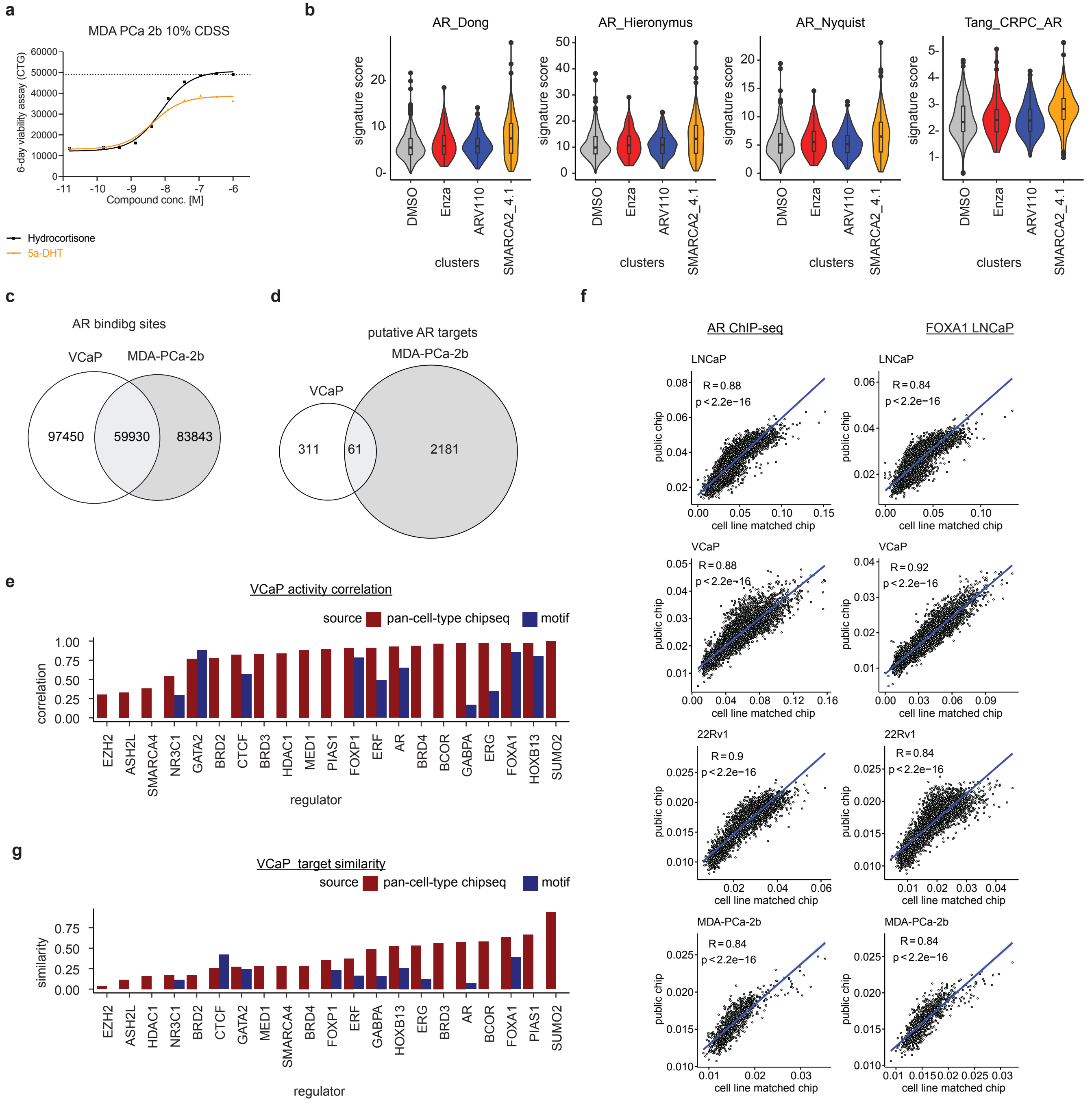
a
MDA PCa 2b 10% CDSS
6-day viability assay (CTG)
60000
50000
40000
30000
20000
10000
0
-11
-10
-9
-8
-7
-6
Compound conc. [M]
Hydrocortisone
5a-DHT
b
AR_Dong
AR_Hieronymus
AR_Nyquist
Tang_CRPC_AR
signature score
clusters
DMSO
Enza
ARV110
SMARCA2_4.1
c
AR bindibg sites
VCaP
MDA-PCa-2b
97450
59930
83843
d
putative AR targets
MDA-PCa-2b
VCaP
311
61
2181
e
VCaP activity correlation
source
pan-cell-type chipseq
motif
correlation
1.00
0.75
0.50
0.25
0.00
EZH2
ASH2L
SMARCA4
NR3C1
GATA2
BRD2
CTCF
BRD3
HDAC1
MED1
PIAS1
FOXP1
ERF
AR
BRD4
BCOR
GABPA
ERG
FOXA1
HOXB13
SUMO2
regulator
f
AR ChIP-seq
FOXA1 LNCaP
LNCaP
R = 0.88
p < 2.2e−16
R = 0.84
p < 2.2e−16
VCaP
R = 0.88
p < 2.2e−16
R = 0.92
p < 2.2e−16
22Rv1
R = 0.9
p < 2.2e−16
R = 0.84
p < 2.2e−16
MDA-PCa-2b
R = 0.84
p < 2.2e−16
R = 0.84
p < 2.2e−16
public chip
cell line matched chip
g
VCaP target similarity
source
pan-cell-type chipseq
motif
similarity
0.75
0.50
0.25
0.00
EZH2
ASH2L
HDAC1
NR3C1
BRD2
CTCF
GATA2
MED1
SMARCA4
BRD4
FOXP1
ERF
GABPA
HOXB13
ERG
BRD3
AR
BCOR
FOXA1
PIAS1
SUMO2
regulator

### Figure S5

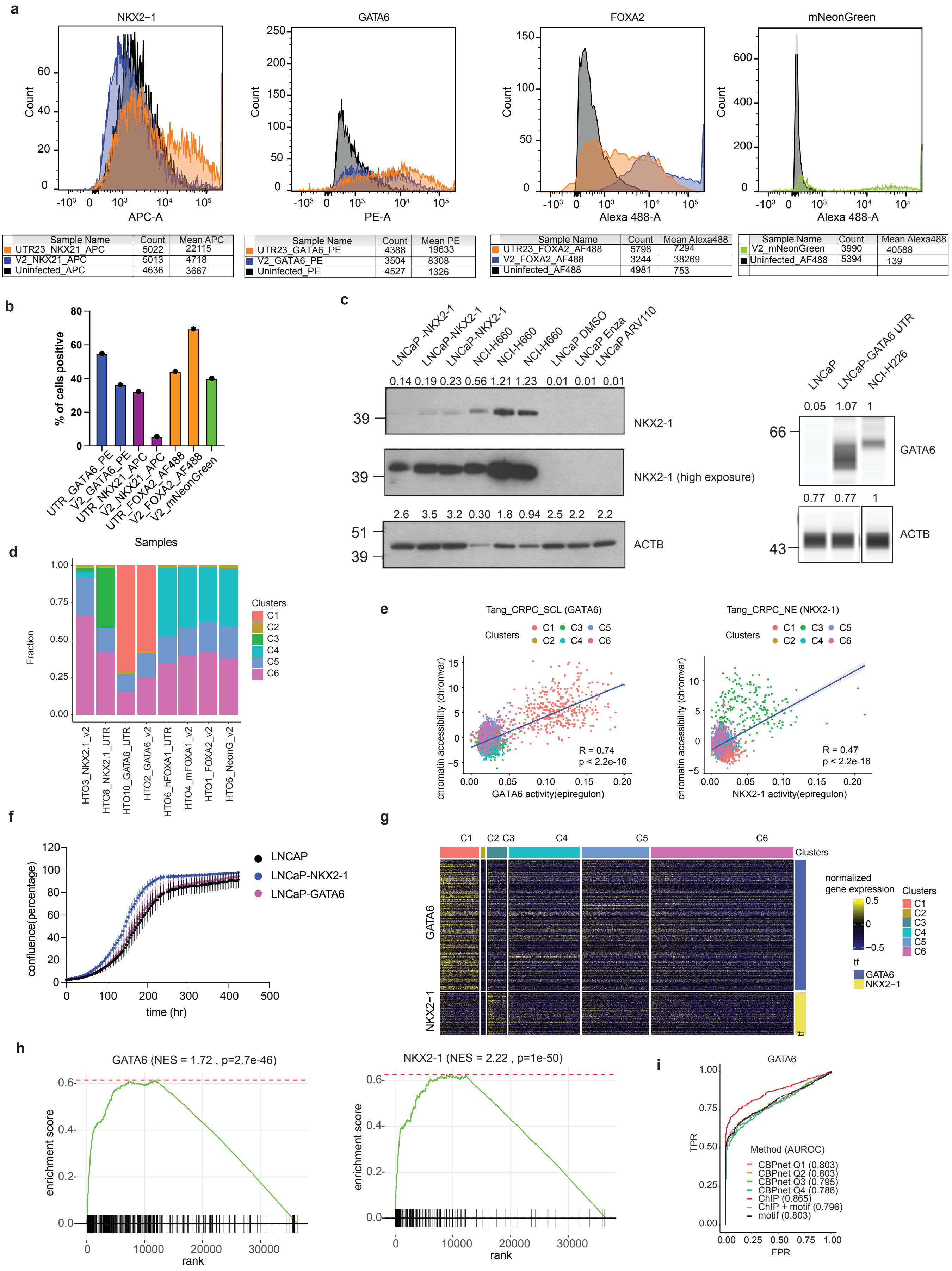


| Sample Name | Count | Mean APC |
|---|---|---|
| UTR23_NKX21_APC | 5022 | 22115 |
| V2_NKX21_APC | 5013 | 4718 |
| Uninfected_APC | 4636 | 3667 |

| Sample Name | Count | Mean PE |
|---|---|---|
| UTR23_GATA6_PE | 4388 | 19633 |
| V2_GATA6_PE | 3504 | 8308 |
| Uninfected_PE | 4527 | 1326 |

| Sample Name | Count | Mean Alexa488 |
|---|---|---|
| UTR23_FOXA2_AF488 | 5798 | 7294 |
| V2_FOXA2_AF488 | 3244 | 38269 |
| Uninfected_AF488 | 4981 | 753 |

| Sample Name | Count | Mean Alexa488 |
|---|---|---|
| V2_mNeonGreen | 3990 | 40588 |
| Uninfected_AF488 | 5394 | 139 |
