## Supplementary Note for "Epiregulon: Inference of single-cell transcription factor activity to predict drug response and drivers of cell state"

**Section 1** Data simulation using scMultiSim to identify default parameters

*Methods*

We used the scMultiSim package[1] to generate gene expression and chromatin accessibility data. This package assigns each cell the cell identity vectors which indicate its internal state. In the discrete mode, the elements of these vectors are sampled from a normal distribution. The differences among the cell identity vectors capture cell-to-cell variability in terms of compound concentrations or morphology. Similarly, different genes are encoded by gene identity vectors. Then vectors of cell states are multiplied by gene identity vectors to produce values used to calculate kinetic parameters for each cell/gene combination. There are 3 kinetic parameters, each one represented by a separate identity vector in the cell or gene: *s* denoting the rate at which mRNA is synthesized, $k_{on}$ indicating the rate at which a gene becomes activated and $k_{off}$ corresponding to the rate of gene inactivation. We used the model agnostic to RNA velocity in which kinetic parameters controlled the number of transcripts according to the following model:

$$p \sim Beta(k_{on}, k_{off})$$

$$n \sim Poisson(p \cdot s)$$

$$y = n\sigma + (1 - \sigma)\frac{k_{on}}{k_{on} + k_{off}}s$$

where *p* is the probability that a gene is active and *n* is the number of transcripts, *y* is the final gene expression, and $\sigma$ is the noise parameter taking the value from 0 to 1 and controlling how much the final expression is affected by the stochasticity imposed by the Beta and Poisson random variables. The final values of the kinetic parameters are sampled from the distribution precomputed with the use of MCMC method. Then both the original and sampled values are being ordered and matched according to their positions.

The simulation was fed with the user-defined gene regulatory network which was used by scMultiSim to create the correct pattern of gene identity vectors. Specifically, target genes inherit from their regulators (transcription factors) the identity vector attributed to the *s* kinetic parameter, which is then multiplied by the connection weight as provided in the GRN input object. The values are then normalized by the number of transcription factors per target gene. The vectors for the regulators are created by randomly choosing two elements, setting their values to 2 and leaving all other elements as zeros. The remaining two types of gene identity vectors, i.e. associated with parameters $k_{on}$ and $k_{off}$, are unaffected by the GRN and sampled from a normal distribution. The cell types were defined by providing as an input to the sim_true_counts function the phylogenetic tree with each cell occupying one out of 9 tree tips. The cell type identity is encoded by the scMultiSim in the respective cell vectors associated with the parameter *s* in two steps procedure: (1) the samples are taken from a multivariate normal distribution in a number corresponding to the length of the cell identity vector and dimensions reflecting the number of cell types with the covariance matrix reflects distances among the cell types on the phylogenetic tree; (2) the values generated in the first step are used as means for the normal distribution from which cell identity vectors are sampled producing variability across cells from the same cluster.

The simulator assigns each target gene 0-2 regulatory regions. No region is shared by any two genes. The values used to determine ATAC-seq signals are calculated by multiplying the cell identity vectors with region identity vectors which are the vectors of random normal variables assigned to each regulatory element. The former are obtained in the same way as vectors associated with the parameter *s*, meaning that they reflect the developmental distances between cell types. The final values for the ATAC-seq are sampled from the distribution that is based on the log-transformed real data. Mapping from simulated to real ATAC-seq values is performed on the basis of the rank match between elements from both sets. The ATAC-seq signal affects the gene expression via the value of the $k_{on}$ parameter of the gene associated with a given region.

We run the simulation on 5000 cells and with UMI protocol. The GRN was prepared in the following way. We set the number of target genes to be 5000 and transcription factors to 500. Then we randomly linked transcription factors to target genes with following constraints: (1) the minimum number of target genes per transcription factor is 3, (2) each target gene should be regulated by at least one transcription factor. The number of target genes per transcription factor was sampled from the Poisson distribution, which was parametrized by an exponential function whose arguments were sampled from uniform distribution over the range [0,5]:

$$n_{targets} \sim Poisson(e^x)$$

where

$$x \sim U(0,5)$$

The weights of the connections between transcription factors and target genes were sampled from the uniform distribution over the range [1,5].

We benchmarked Epiregulon in various simulation conditions. Besides generating data as described above, we also used the add_expr_noise function to make the result more realistic by adding variations from amplification bias, dropouts, differences in capture efficiency during library preparation etc. We tested three levels of technical noise: one with default values of add_expr_noise parameters (alpha_mean = 0.1, alpha_sd = 0.02, alpha_gene_mean = 1, alpha_gene_sd = 0, depth_mean = 1e5, depth_sd = 3e3, atac.obs.prob = 0.3, atac.sd.frac = 0.5), one with stronger (alpha_mean = 0.05, alpha_sd = 0.005, alpha_gene_mean = 0.75, alpha_gene_sd = 0.05, depth_mean = 5e4, depth_sd = 2e3, atac.obs.prob = 0.2, atac.sd.frac = 0.2) noise and one with weaker noise (alpha_mean = 0.5, alpha_sd = 0.001, alpha_gene_mean = 1, alpha_gene_sd = 0, depth_mean = 2e5, depth_sd = 1e3, atac.obs.prob = 0.6, atac.sd.frac = 0.05). These parameters control for the mean cell and gene specific capture efficiency, sequence, and ATAC-seq signal sparsity and variation of these parameters across the cells and genes. Moreover, we run a simulation with two different values for the atac.signal parameter, default (0.5) and greater (0.85). It controls how much $k_{on}$ parameters depend on the ATAC-seq signal.

To calculate the true transcription factor activity, we took the sum of differences in its target genes expression generated from either the original GRN or from a GRN obtained by ablating the activity of a transcription factor through setting its edge weights to close to zero. We repeat this process for each of the 500 transcription factors. Using the same generator seed, we ensured that the additional simulated gene expression and chromatin accessibility matrices were matched to the original matrices by cells. We set parameter controlling intrinsic noise to 0 to ensure that the only source of variation in the results are the differences in GRN.

Then we compared the activity scores with those outputted from the workflow offered by the Epiregulon package. Simulated gene expression and ATAC-seq data were used to create SingleCellExperiment objects with cell type identity provided in colData slot. The gene expression was normalized to the total number of $10^4$ counts per cell and subject to PCA whose results were also included in the SingleCellExperiment object. We tested all available methods for calculation of GRN weights i.e. Wilcoxon, corr and MI. For corr and MI, cell types were used as the cluster argument to *addWeights* function. Moreover, we compared the Epiregulon performance depending on whether cell aggregation has been done by setting aggregateCells parameter to TRUE or FALSE. We tested separately the performance of the pruneRegulon function across 3 values of regulon_cutoff argument (0.5, 0.2, 0.05, 0.01) by contamination of GRN with false connections. Apart from this case we skipped network pruning in Epiregulon workflow since here the simulation was run on the uncontaminated GRN.

*Results*

We used simulations to understand the choice of weight estimation methods and the effects of false GRN connections, data sparsity and the contribution of RE accessibility on the accuracy of TF activity inference by Epiregulon. We generated simulated single-cell multiomics data from a ground truth GRN using the scMultiSim package[1]. Each cell was characterized by vectors of cell identity parameters which were then used to create gene expression data and ATAC-seq signal (Supplementary Note Figure 1a). We perturbed each TF by setting the weights to 0 and calculated a ground truth TF activity based on the effects of the perturbation on the target gene expression. This models the situation in which the TF is still being transcribed but has lost its regulatory potential, e.g., due to chemical inhibition of the TF protein itself. We utilized cell identity to pair the outputs of the basic and perturbed simulation. We quantified the performance by correlating estimated TF activity against true activity.

For all the methods, we consider both the TF expression and RE accessibility when estimating the weights. Amongst the three weight methods, mutual information was the best performing method but the differences between the different methods started to diminish with increasing data sparsity (Supplementary Note Figure 1b). When we introduced false connections, the performance of the mutual information method started to deteriorate and the different methods performed similarly (Supplementary Note Figure 1c). Aggregation of similar cells further improved the performance of the Wilcoxon method (Supplementary Note Figure 1d). All weight methods saw an improvement in their performance when gene expression was increasingly dependent on chromatin accessibility (Supplementary Note Figure 1e). Based on a GRN consisting of 50% false connections and 50% true connections, we showed that pruning increases the composition of true connections albeit at the expense of losing true targets (Supplementary Note Figure 1f). Even in the absence of pruning, the weight estimation already buffered against GRN contamination by assigning higher weights to true connections than to false ones (Supplementary Note Figure 1g). Therefore, the presence of false connections does not profoundly impact the final TF activity (Supplementary Note Figure 1h). However, given that the size of a regulon averages to more than

100 targets per TF in a real dataset, we reasoned that removing false connections will result in a cleaner set of target genes for downstream analysis such as gene set enrichment and Jaccard similarity calculations. Taken together, simulated data suggests that all methods (Wilcoxon, correlation and mutual information) perform similarly and robustly in the presence of data sparsity and false connections and pruning can enhance the composition of true GRN connections.

### Supplementary Note Figure 1

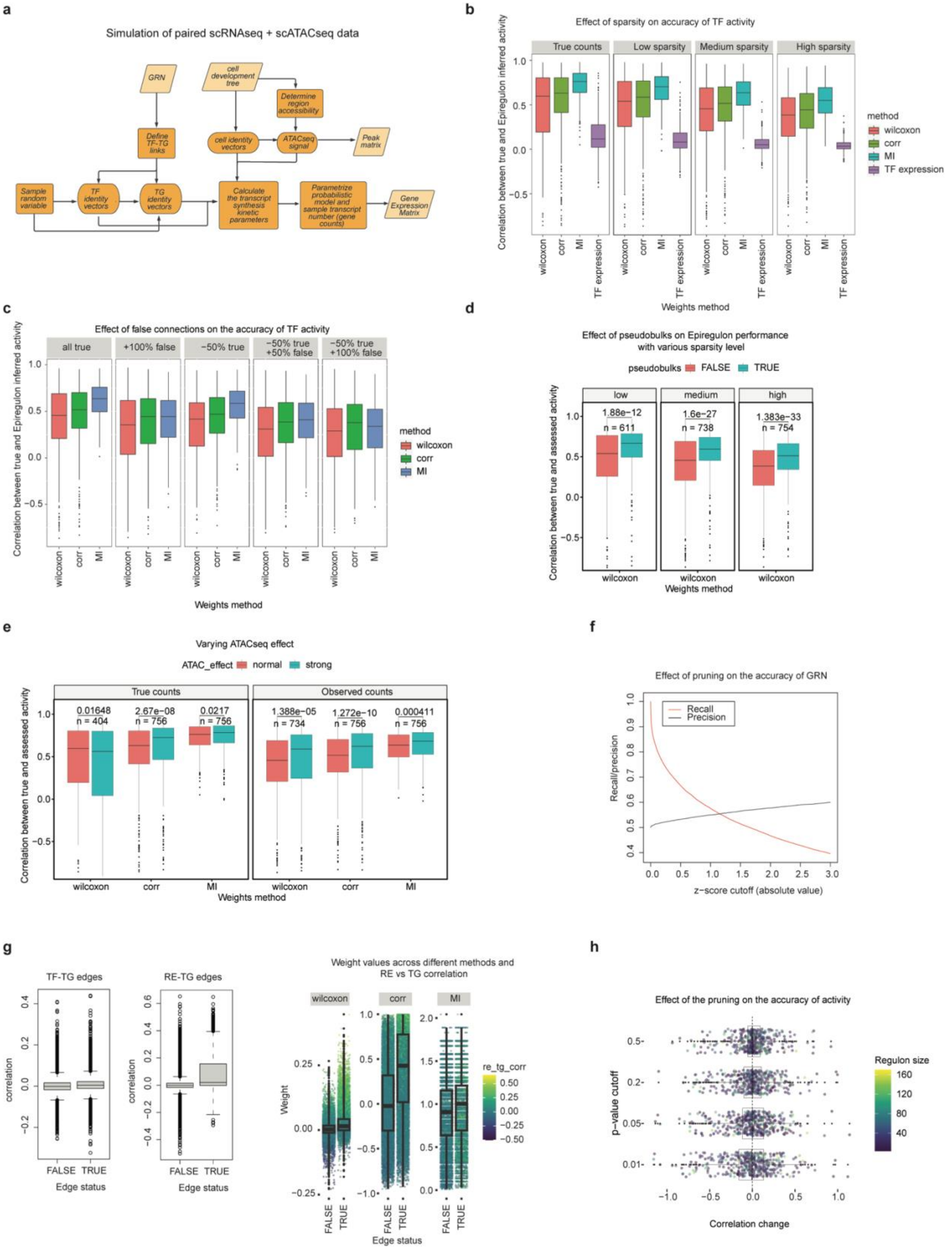

**Supplementary Note Figure 1 Data simulation using scMultiSim**

a) We used the scMultiSim package to simulate paired scRNA-seq and scATAC-seq data. Briefly, we provided a lineage tree that specified the identities of the individual cells and a ground truth GRN that specified the regulatory relationships between TF-TG and RE-TG. Gene expression and peak matrices were generated taking into account both cell identities and gene regulatory relationships. We obtained the ground truth activity of each TF by summing the differences in target gene expression as defined by the GRN. For all assessments from b-e, we use the fixed cutoff and tf_re.merge set to TRUE.
b) We varied the degree of sparsity by increasing the technical noise and evaluated the correlation of activity against ground truth TF activity across all genes for each of the weight methods. Each data point represents a TF. The ATAC effect was set at basic. GRN had 100% true connections.
c) We added noise to GRN by adding false connections and removing true connections, and correlated the activity against ground truth activity for each TF. The ATAC effect was set at basic and data sparsity was set at medium.
d) Cell clusters were first defined using k-means clustering with a median cluster size of 10 cells for the Wilcoxon method. We correlated the activity against ground truth activity for each TF. The ATAC effect was set at normal and data sparsity was set at medium. GRN had 100% true connections. n indicates the number of TFs retrieved in the benchmarking.
e) We varied the effect of ATAC on the gene expression, with the strong ATAC effect indicating a greater importance of ATAC counts on shaping gene expression counts. GRN had 100% true connections. True counts were generated without data sparsity and observed counts were generated with medium data sparsity. n indicates the number of TFs retrieved in the benchmarking.
f) We spiked in 100% false connections to the true GRN. We examined the recall and precision of true GRN connections with varying degrees of z score cutoffs computed by chi-square statistics
g) We spiked in 100% false connections to the true GRN. We show that true connections have increased RE-TG correlations and this translated to higher weights in the true connections.
h) We evaluated the effects of pruning on the accuracy of TF activity predictions. We computed the ground truth TF activity from the predefined GRN with high data sparsity and strong ATAC effects. We spiked in 100% false connections and compared the correlation between the TF activity inferred from either the unpruned or pruned GRN vs the ground truth GRN. Shown is the difference between the two correlations. A positive difference indicates improved performance due to pruning.

**Section 2** Data simulation of differential network

*Methods*

We used simulations to evaluate two algorithms for performing pairwise differential network analysis. These simulations were designed to characterize each algorithm's performance at detecting transcription factors (TFs) whose local network connectivity with regulatory elements (REs) and target genes (TGs) differed in several simple scenarios in which the true signal was known.

To do so, we first simulated a "ground truth" network comprising 250 TFs, 1000 REs, and 500 TGs. In this ground truth network, each TF-RE pair was randomly assigned a scalar value (distributed along a standard normal) to represent the true regulatory association between the TF and the RE. Similarly, each RE-TG pair was also assigned scalar values according to the same procedure to represent the true regulatory association between each RE and each TG. These scalar values were stored as a [250 x 1000] TF-RE adjacency matrix U and a [1000 x 500] adjacency matrix V.

To simulate the sparsity present in real-world gene regulatory networks, entries in both U and V were randomly dropped out to 0 with a random probability between 0.2 and 0.8 that was assigned to each TF independently in order to provide a set of TFs and REs with a range of total regulatory activity (represented by their total number of nonzero edges in the U and V matrices, respectively).

Once this ground truth network was created, it was used to simulate pairs of networks (representing different cell types or experimental conditions) with specific differences – the goal of our differential network analysis algorithms would be to retrieve these differences with high fidelity. Specifically, the ground truth network was mutated in the following ways to generate differential networks representing distinct cell types:

1) **The "reference" (default) cell type**: Noise was added to the base network by randomly permuting some percentage of edges in both U and V randomly, thus maintaining the same degree of sparsity as the ground truth network, but with additional noise (simulating an estimate of the true ground truth network that would come from using real-world data)
2) **The edge-enriched cell type:** Additional edges were added between a user-specified subset of the TFs and a random subset of the REs in the ground truth network. The number of edges added is given as a percentage of each TF's number of nonzero edges in the ground truth network (to maintain differences in TF-RE connectivity between TFs). Additional noise is also added by permuting some subset of the nonzero edges as above, to simulate the estimation process from real-world data. We call this type of signal **edge enrichment**, and it represents the biological scenario in which a TF's regulatory activity increases in one cell type compared to another.
3) **The edge-depleted cell type:** Edges are removed between a user-specified subset of the TFs and a random subset of the REs in the ground truth network. The number of edges removed is given as a percentage of each TF's number of nonzero edges in the base network (to maintain differences in TF-RE connectivity between TFs). Additional noise is also added by permuting some subset of the nonzero edges as above, to simulate the estimation process

from real-world data. We call this type of signal **edge depletion**, and it represents the biological scenario in which a TF's regulatory activity decreases in one cell type compared to another.

4) **The edge-permuted cell type:** Edges are permuted within a user-specified subset of the TFs such that the number of TF-RE edges for each TF remains the same, but with different TF-RE connectivity. The number of edges removed is given as a percentage of each TF's number of nonzero edges in the base network (to maintain differences in TF-RE connectivity between TFs). Additional noise is also added by permuting some subset of the nonzero edges as above, to simulate the estimation process from real-world data. We call this type of signal **edge permutation**, and it represents the biological scenario in which a TFs overall regulatory activity neither increases nor decreases overall, but rather shifts from one regulatory program to another.

Using these cell-type-specific U and V matrices, we construct TF-TG bipartite graphs in which each TF-TG pair is connected by either a single edge (if only one RE connects the TF and TG) or multiple edges (if multiple REs connect the TF and TG). The weight of each edge in the TF-TG graph is the product of the entry in the U matrix corresponding to the TF-RE edge and the entry in the V matrix corresponding to the RE-TG edge.

Finally, we can use these cell type-specific graphs to rank the TFs from most differential between cell types to least differential between cell types using either of 2 algorithms. The first algorithm (**centrality subtraction**) first calculates the degree centrality of each TF in each cell-type specific graph. Then, the degree centrality values for each TF are subtracted between the pair of graphs being compared, and the TFs are ranked in order of the highest to lowest absolute value of the degree centrality difference. The second algorithm (**edge subtraction**) first calculates the difference in the weight of each TF-TG edge in each cell-type specific graph. Then, the degree centrality values for each TF in the edge-subtracted graph are calculated, and the TFs are ranked in order of the highest to lowest degree centrality in the edge-subtracted graph. Using these ranked lists of TFs, we calculate ROC curves for each approach (by varying the threshold used to separate differential vs. non-differential TFs.

Under this general framework, we evaluated how both the centrality subtraction and edge subtraction algorithms performed as we varied the amount of noise used in network "estimation" (i.e. the percentage of permuted edges in the ground truth network) as well as the amount of signal used to distinguish differential TFs and non-differential TFs in each cell type (edge-enriched, edge-depleted, and edge-permuted). Specifically, in experiments varying the amount of noise, the signal (i.e. proportion of enriched, depleted, or permuted edges) was fixed at 0.15 and varying the proportion of permuted edges for each TF's edges in the ground truth network 0 to 0.5. In experiments varying the amount of signal, we fixed the noise at a permutation of 0.1 of each TF's edges from the ground truth network and varied the proportion of enriched, depleted, or permuted edges from 0 to 0.25.

*Results*

Using simulated data, we demonstrated that edge subtraction outperforms centrality subtraction. Both methods perform comparably in detecting edge enrichment and depletion, but edge subtraction excels in detecting edge permutation (see Supplementary Note Figure 2a-b). The edge subtraction method proves to be both robust and sensitive, as it can discern differences between networks even in the presence of considerable noise (as depicted in Supplementary Note Figure 2a) and at relatively low levels of signal (as shown in Supplementary Note Figure 2b).

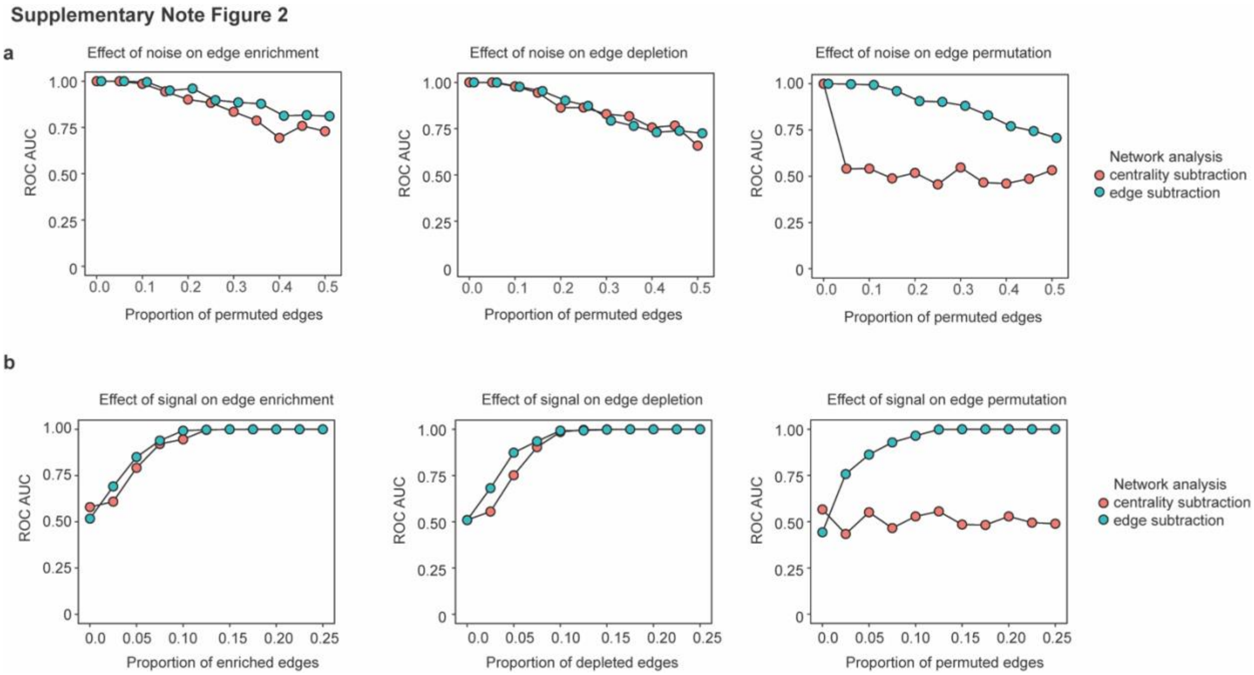


**Supplementary Note Figure 2: Network simulation**

a) We added noise to the network by increasing proportions of randomly permuted edges on top of true differential edges (15% of total edges). Centrality subtraction was computed by subtracting the degree centralities of two networks whereas edge subtraction was computed by centrality of the edge subtracted network. See methods for details.
b) We varied the signal strength by increasing the proportions of altered edges (enrichment, depletion or permutations) with noise fixed at 10% of total edges.

1 Li, H., Zhang, Z., Squires, M., Chen, X. & Zhang, X. scMultiSim: simulation of multi-modality single cell data guided by cell-cell interactions and gene regulatory networks. *bioRxiv*, 2022.2010.2015.512320, doi:10.1101/2022.10.15.512320 (2022).
